# Functional reorganization of brain networks across the human menstrual cycle

**DOI:** 10.1101/866913

**Authors:** Laura Pritschet, Tyler Santander, Caitlin M. Taylor, Evan Layher, Shuying Yu, Michael B. Miller, Scott T. Grafton, Emily G. Jacobs

**Author notes:** Correspondence: Emily G. Jacobs, Department of Psychological & Brain Sciences, University of California, Santa Barbara, Santa Barbara, CA 93106. Authors contributed equally to this work (Laura Pritschet, Tyler Santander).

## Abstract

The brain is an endocrine organ, sensitive to the rhythmic changes in sex hormone production that occurs in most mammalian species. In rodents and nonhuman primates, estrogen and progesterone’s impact on the brain is evident across a range of spatiotemporal scales. Yet, the influence of sex hormones on the functional architecture of the human brain is largely unknown. In this dense-sampling, deep phenotyping study, we examine the extent to which endogenous fluctuations in sex hormones alter intrinsic brain networks at rest in a woman who underwent brain imaging and venipuncture for 30 consecutive days. Standardized regression analyses illustrate estrogen and progesterone’s widespread associations with functional connectivity. Time-lagged analyses examined the temporal directionality of these relationships and suggest that cortical network dynamics (particularly in the Default Mode and Dorsal Attention Networks, whose hubs are densely populated with estrogen receptors) are preceded—and perhaps driven—by hormonal fluctuations. A similar pattern of associations was observed in a follow-up study one year later. Together, these results reveal the rhythmic nature in which brain networks reorganize across the human menstrual cycle. Neuroimaging studies that densely sample the individual connectome have begun to transform our understanding of the brain’s functional organization. As these results indicate, taking endocrine factors into account is critical for fully understanding the intrinsic dynamics of the human brain.

**Highlights:**

- Intrinsic fluctuations in sex hormones shape the brain’s functional architecture.
- Estradiol facilitates tighter coherence within whole-brain functional networks.
- Progesterone has the opposite, reductive effect.
- Ovulation (via estradiol) modulates variation in topological network states.
- Effects are pronounced in network hubs densely populated with estrogen receptors.

## 1 Introduction

The brain is an endocrine organ whose day-to-day function is intimately tied to the action of neuromodulatory hormones (Frick et al., 2015; Galea et al., 2017; Hara et al., 2015; Woolley and McEwen, 1993). Yet, the study of brain-hormone interactions in human neuroscience has often been woefully myopic in scope: the classical approach of interrogating the brain involves collecting data at a single time point from multiple subjects and averaging across individuals to provide evidence for a hormone-brain-behavior relationship. This cross-sectional approach obscures the rich, rhythmic nature of endogenous hormone production. A promising trend in network neuroscience is to flip the cross-sectional model by tracking small samples of individuals over timescales of weeks, months, or years to provide insight into how biological, behavioral, and state-dependent factors influence intra- and inter-individual variability in the brain’s intrinsic network organization (Gordon et al., 2017; Gratton et al., 2018a; Poldrack et al., 2015). Neuroimaging studies that densely sample the individual connectome are beginning to transform our understanding of the dynamics of human brain organization. However, these studies commonly overlook sex steroid hormones as a source of variability—a surprising omission given that sex hormones are powerful neuromodulators that display stable circadian, infradian, and circannual rhythms in nearly all mammalian species. In the present study, we illustrate robust, time-dependent interactions between the sex steroid hormones 17*β*-estradiol and progesterone during a complete menstrual cycle. A within-subject replication study further confirms the robustness of these effects. These results offer compelling evidence that sex hormones modulate widespread patterns of connectivity in the human brain.

Converging evidence from rodent (Frick et al., 2018, 2015; Woolley and McEwen, 1993), non-human primate (Hao et al., 2006; Wang et al., 2010), and human neuroimaging studies (Berman et al., 1997; Jacobs and D’Esposito, 2011; Jacobs et al., 2016a,b; Lisofsky et al., 2015; Petersen et al., 2014) has established the widespread influence of 17*β*-estradiol and progesterone on regions of the mammalian brain that support higher level cognitive functions. Estradiol and progesterone signaling are critical components of cell survival and plasticity, exerting excitatory and inhibitory effects that are evident across multiple spatial and temporal scales (Frick et al., 2018; Galea et al., 2017). The dense expression of estrogen and progesterone receptors (ER; PR) in cortical and subcortical tissue underscores the widespread nature of hormone action. For example, in non-human primates, ∼50% of pyramidal neurons in prefrontal cortex (PFC) express ER (Wang et al., 2010) and estradiol regulates dendritic spine proliferation in this region (Hara et al., 2015). Across the rodent estrous cycle (occurring every 4-5 days), fluctuations in estradiol enhance spinogenesis in hippocampal CA1 neurons, while progesterone inhibits this effect (Woolley and McEwen, 1993).

During an average human menstrual cycle, occurring every 25-32 days, women experience a ∼12-fold increase in estradiol and an ∼800-fold increase in progesterone. Despite this striking change in endocrine status, we lack a complete understanding of how the large-scale functional architecture of the human brain responds to rhythmic changes in sex hormone production across the menstrual cycle. Much of our understanding of cycle-dependent changes in brain structure (Sheppard et al., 2019; Woolley and McEwen, 1993) and function (Hampson et al., 2014; Kim and Frick, 2017; Warren and Juraska, 1997) comes from rodent studies, since the length of the human menstrual cycle (at least 5× longer than rodents’ estrous cycle) presents experimental hurdles that make longitudinal studies challenging. A common solution is to study women a few times throughout their cycle, targeting stages that roughly correspond to peak/trough hormone concentrations. Using this ‘sparse-sampling’ approach, studies have examined resting-state connectivity in discrete stages of the cycle (De Bondt et al., 2015; Hjelmervik et al., 2014; Lisofsky et al., 2015; Petersen et al., 2014; Syan et al., 2017; Weis et al., 2019); however, some of these findings are undermined by inconsistencies in cycle staging methods, lack of direct hormone assessments, or limitations in functional connectivity methods.

In this dense-sampling, deep-phenotyping study, we determined whether day-to-day variation in sex hormone concentrations impacts connectivity states across major intrinsic brain networks. First, we assessed brain-hormone interactions over 30-consecutive days representing a complete menstrual cycle (Study 1). To probe the reliability of these findings, procedures were then repeated over a second 30-day period, providing a within-subject controlled replication (Study 2). Results reveal that intrinsic functional connectivity is linearly dependent on hormonal dynamics across the menstrual cycle at multiple spatiotemporal scales. Estradiol and progesterone were associated with spatially-diffuse changes in connectivity, both at time-synchronous and time-lagged levels of analysis, demonstrating that intrinsic fluctuations in sex hormones—particularly the ovulatory surge in estradiol—may contribute to dynamic variation in the functional network architecture of the human brain. We further highlight this sensitivity to estradiol in a controlled replication study. Together, these findings provide insight into how brain networks reorganize across the human menstrual cycle and suggest that consideration of the hormonal milieu is critical for fully understanding the intrinsic dynamics of the human brain.

## 2 Materials and Methods

### 2.1 Participants

The participant (author L.P.) is a right-handed Caucasian female, aged 23 years for duration of the study. The participant had no history of neuropsychiatric diagnosis, endocrine disorders, or prior head trauma. She had a history of regular menstrual cycles (no missed periods, cycle occurring every 26-28 days) and had not taken hormone-based medication in the 12 months prior to the first study. The participant gave written informed consent and the study was approved by the University of California, Santa Barbara Human Subjects Committee.

### 2.2 Study design

The participant underwent testing for 30 consecutive days, with the first test session determined independently of cycle stage for maximal blindness to hormone status (Study 1). One year later, as part of a larger parent project, the participant repeated the 30-day protocol while on a hormone regimen (0.02mg ethinyl-estradiol, 0.1mg levonorgestrel, Aubra, Afaxys Pharmaceuticals), which she began 10 months prior to the start of data collection (Study 2). The general procedures for both studies were identical (**Figure 1**). The pharmacological regimen used in Study 2 chronically and selectively suppressed progesterone while leaving estradiol dynamics largely indistinguishable from Study 1. This provided a natural replication dataset in which to test the reliability of the estradiol associations observed in the first study. The participant began each test session with a daily questionnaire (see **Section 2.3: Behavioral assessments**), followed by an immersive reality spatial navigation task (not reported here). Time-locked collection of serum and whole blood started each day at 10:00am in Study 1 and 11:00am in Study 2 (*±* 30 min), when the participant gave a blood sample. Endocrine samples were collected, at minimum, after two hours of no food or drink consumption (excluding water). The participant refrained from consuming caffeinated beverages before each test session. The MRI session lasted one hour and consisted of structural and functional MRI sequences.

**Figure 1.**
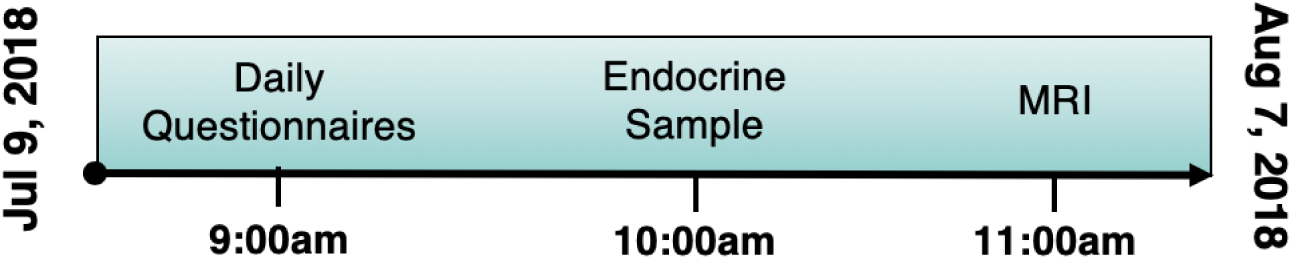
Timeline of data collection for the 30 experiment sessions. Endocrine and MRI assessments were collected at the same time each day to minimize time-of-day effects.

### 2.3 Behavioral assessments

To monitor state-dependent mood and lifestyle measures over the cycle, the following scales (adapted to reflect the past 24 hours) were administered each morning: Perceived Stress Scale (PSS) (Cohen et al., 1983), Pittsburgh Sleep Quality Index (PSQI) (Buysse et al., 1989), State-Trait Anxiety Inventory for Adults (STAI) (Spielberger and Vagg, 1984), and Profile of Mood States (POMS) (Pollock et al., 1979). The participant had moderate levels of anxiety as determined by STAI reference ranges; however, all other measures fell within the ‘normal’ standard range. Self-reported stress was marginally higher in Study 2 (*M*_*diff*_ = 3.9, *t*(58) = 2.66, *p* = .046); no other differences in mood or lifestyle measures were observed between the two studies. Few significant relationships were observed between hormones and state-dependent measures following FDR-correction for multiple comparisons (*q* < .05)—and critically, none of these state-dependent factors were associated with estradiol (**Figure 2A**). Furthermore, performance on a daily selective attention task (Cohen et al., 2014) was stable across the experiment (*M* = 98%, *SD* = 0.01; **Figure 2B**). Taken together, there were no indications of significant shifts in behavior across the cycle.

**Figure 2.**
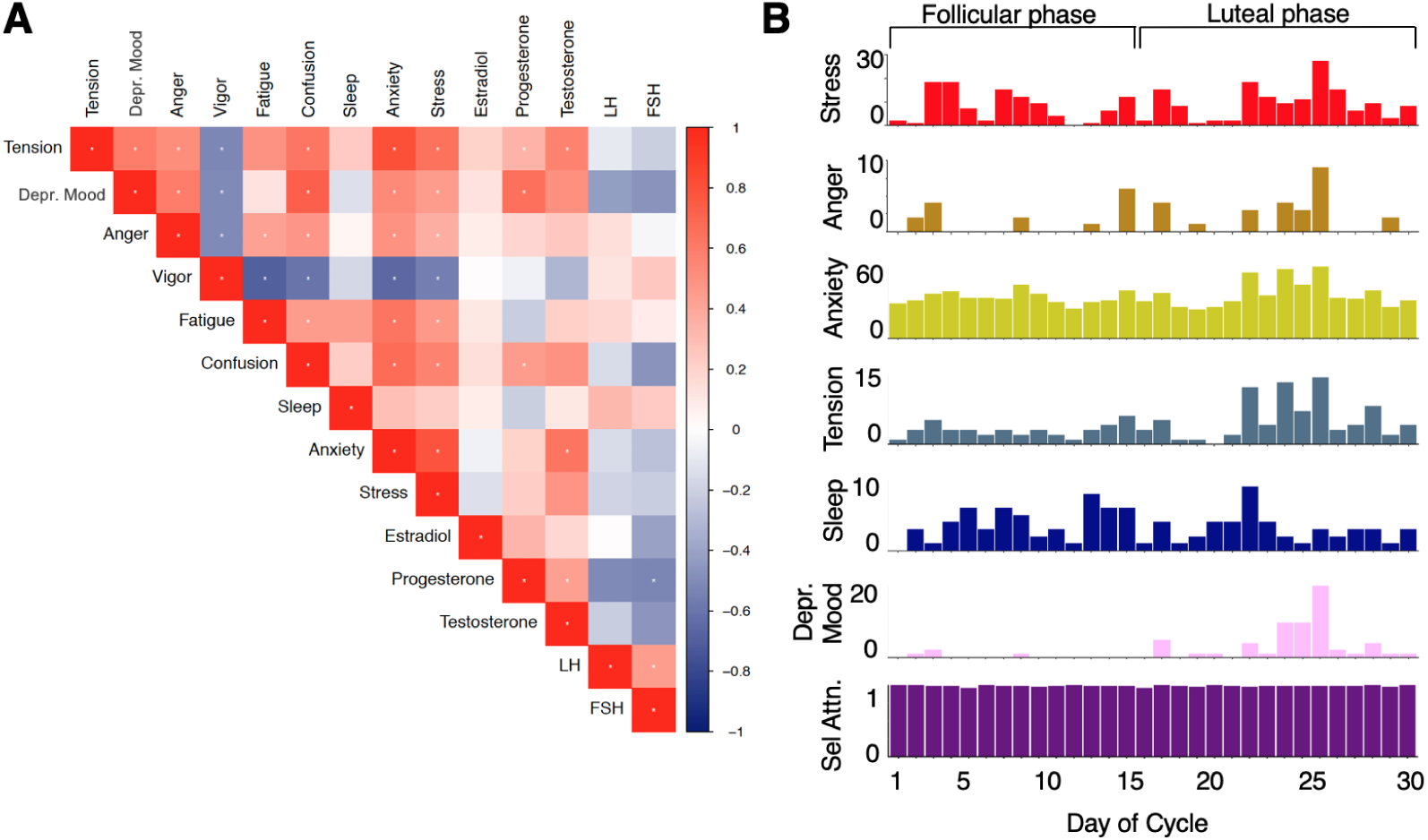
Behavioral variation across the 30-day experiment (Study 1). (**A**) Correlation plot showing relationships between mood, lifestyle measures, and sex steroid hormone concentrations. Cooler cells indicate negative correlations, warm cells indicate positive correlations, and white cells indicate no relationship. Asterisks indicate significant correlations after FDR-correction (*q* < .05). (**B**) Mood and lifestyle measures vary across the cycle; cognitive performance (selective attention) does not. ‘Day 1’ indicates first day of menstruation, *not* first day of experiment. Abbreviations: LH, Lutenizing hormone; FSH, Follicle-stimulating hormone.

### 2.4 Endocrine procedures

A licensed phlebotomist inserted a saline-lock intravenous line into the dominant or non-dominant hand or forearm daily to evaluate hypothalamic-pituitary-gonadal axis hormones, including serum levels of gonadal hormones (17*β*-estradiol, progesterone and testosterone) and the pituitary gonadotropins luteinizing hormone (LH) and follicle stimulating hormone (FSH). One 10cc mL blood sample was collected in a vacutainer SST (BD Diagnostic Systems) each session. The sample clotted at room temperature for 45 min until centrifugation (2,000 ×*g* for 10 minutes) and were then aliquoted into three 1 mL microtubes. Serum samples were stored at -20°C until assayed. Serum concentrations were determined via liquid chromatography-mass spectrometry (for all steroid hormones) and immunoassay (for all gonadotropins) at the Brigham and Women’s Hospital Research Assay Core. Assay sensitivities, dynamic range, and intra-assay coefficients of variation (respectively) were as follows: estradiol, 1 pg/mL, 1–500 pg/mL, < 5% relative standard deviation (*RSD*); progesterone, 0.05 ng/mL, 0.05–10 ng/mL, 9.33% *RSD*; testosterone, 1.0 ng/dL, 1–2000 ng/dL, < 4% *RSD*. FSH and LH levels were determined via chemiluminescent assay (Beckman Coulter). The assay sensitivity, dynamic range, and the intra-assay coefficient of variation were as follows: FSH, 0.2 mIU/mL, 0.2–200 mIU/mL, 3.1–4.3%; LH, 0.2 mIU/mL, 0.2–250 mIU/mL, 4.3–6.4%. Importantly, we note that LC-MS assessments of *exogenous* hormone concentrations (attributable to the hormone regimen itself) showed that serum concentrations of ethinyl estradiol were very low (*M* = 0.01 ng/mL; range: 0.001–0.016 ng/mL) and below 1.5 ng/mL for levonorgestrel (*M* = 0.91 ng/mL; range: 0.03–1.43 ng/mL): this ensures that the brain-hormone associations reported below are still due to *endogenous* estradiol action in Study 2.

### 2.5 MRI acquisition

The participant underwent a daily magnetic resonance imaging scan on a Siemens 3T Prisma scanner equipped with a 64-channel phased-array head coil. First, high-resolution anatomical scans were acquired using a *T*_1_-weighted magnetization prepared rapid gradient echo (MPRAGE) sequence (TR = 2500 ms, TE = 2.31 ms, TI = 934 ms, flip angle = 7°; 0.8 mm thickness) followed by a gradient echo fieldmap (TR = 758 ms, TE_1_ = 4.92 ms, TE_2_ = 7.38 ms, flip angle = 60°). Next, the participant completed a 10-minute resting-state fMRI scan using a *T*_2_^*^-weighted multiband echo-planar imaging (EPI) sequence sensitive to the blood oxygenation level-dependent (BOLD) contrast (TR = 720 ms, TE = 37 ms, flip angle = 56°, multiband factor = 8; 72 oblique slices, voxel size = 2 mm^3^). In an effort to minimize motion, the head was secured with a custom, 3D-printed foam head case (https://caseforge.co/) (days 8–30 of Study 1, days 1–30 of Study 2). Overall motion (mean framewise displacement) was negligible (**Figure S1**), with fewer than 130 microns of motion on average each day. Importantly, mean framewise displacement was also not correlated with estradiol concentrations (Study 1: Spearman *r* = −0.06, *p* = .758; Study 2: Spearman *r* = −0.33, *p* = .071).

### 2.6 fMRI preprocessing

Initial preprocessing was performed using the Statistical Parametric Mapping 12 software (SPM12, Wellcome Trust Centre for Neuroimaging, London) in Matlab. Functional data were realigned and unwarped to correct for head motion and geometric deformations due to motion and magnetic field inhomogeneities; the mean motion-corrected image was then coregistered to the high-resolution anatomical image. All scans were then registered to a subject-specific anatomical template created using Advanced Normalization Tools’ (ANTs) multivariate template construction (**Figure S2**). A 4 mm full-width at half-maximum (FWHM) isotropic Gaussian kernel was subsequently applied to smooth the functional data. Further preparation for resting-state functional connectivity was implemented using in-house Matlab scripts. Global signal scaling (median = 1,000) was applied to account for transient fluctuations in signal intensity across space and time, and voxelwise timeseries were linearly detrended. Residual BOLD signal from each voxel was extracted after removing the effects of head motion and five physiological noise components (CSF + white matter signal). Motion was modeled based on the Friston-24 approach, using a Volterra expansion of translational/rotational motion parameters, accounting for autoregressive and nonlinear effects of head motion on the BOLD signal (Friston et al., 1996). All nuisance regressors were detrended to match the BOLD timeseries. Our use of coherence allows for the estimation of frequency-specific covariances in spectral components below the range contaminated by physiological noise. Nevertheless, to ensure the robustness of our results, we re-analyzed the data with global signal regression included. This had little bearing on the overall findings. For completeness, results from the GSR-based processing pipeline are provided in the **Supplementary Material**.

### 2.7 Functional connectivity estimation

Functional network nodes were defined based on a 400-region cortical parcellation (Schaefer et al., 2018) and 15 regions from the Harvard-Oxford subcortical atlas (http://www.fmrib.ox.ac.uk/fsl/). For each day, a summary timecourse was extracted per node by taking the first eigenvariate across functional volumes (Friston et al., 2006). These regional timeseries were then decomposed into several frequency bands using a maximal overlap discrete wavelet transform (Daubechies extremal phase filter, length = 8). Low-frequency fluctuations in wavelets 3–6 (∼0.01–0.17 Hz) were selected for subsequent connectivity analyses (Patel and Bullmore, 2016). We estimated the *spectral* association between regional timeseries using magnitude-squared coherence: this yielded a 415 × 415 functional association matrix each day, whose elements indicated the strength of functional connectivity between all pairs of nodes (FDR-thresholded at *q* < .05). Coherence offers several advantages over alternative methods for assessing connectivity: 1) estimation of *frequency-specific covariances*, 2) *simple interpretability* (values are normalized to the [0, 1] interval), and 3) *robustness to temporal variability in hemodynamics* between brain regions, which can otherwise introduce time-lag confounds to connectivity estimates via Pearson correlation.

### 2.8 Statistical analysis

First, we assessed time-synchronous variation in functional connectivity associated with estradiol and progesterone through a standardized regression analysis. Data were *Z*-transformed and edgewise coherence was regressed against hormonal timeseries to capture day-by-day variation in connectivity relative to hormonal fluctuations. For each model, we computed robust empirical null distributions of test-statistics (*β/SE*) via 10,000 iterations of nonparametric permutation testing: under the null hypothesis of no temporal association between connectivity and hormones, the coherence data at each edge were randomly permuted, models were fit, and two-tailed *p*-values were obtained as the proportion of models in which the absolute value of the permuted test statistics equaled or exceeded the absolute value of the ‘true’ test statistics. We report edges surviving a threshold of *p* < .001. We did not apply additional corrections in an effort to maximize power in our small sample size; Study 2 instead offers an independent validation of the observed whole-brain effects.

Next, we sought to capture linear dependencies between hormones and network connectivity *directed in time* using vector autoregressive (VAR) models. Here we chose to focus exclusively on estradiol for two reasons: 1) the highly-bimodal time-course of progesterone over a natural cycle confers a considerably longer autocorrelative structure, requiring many more free parameters (i.e. higher-order models, ultimately affording fewer degrees of freedom); and 2) progesterone lacks an appreciable pattern of periodicity in its autocovariance with network timeseries, suggesting less relevance for time-lagged analysis over a single cycle. In contrast, estradiol has a much smoother time-course that is well-suited for temporal evolution models such as VAR.

In short, VAR solves a simultaneous system of equations that fits *current* states of the brain and estradiol from the *previous* states of each. For consistency, we considered only *second-order* VAR models, given a fairly reliable first zero-crossing of brain/hormone autocovariance functions at lag two (this was based on common criteria noted in other instances of time-delayed models; Boker et al. (2014)). Fit parameters for each VAR therefore reflect the following general form:

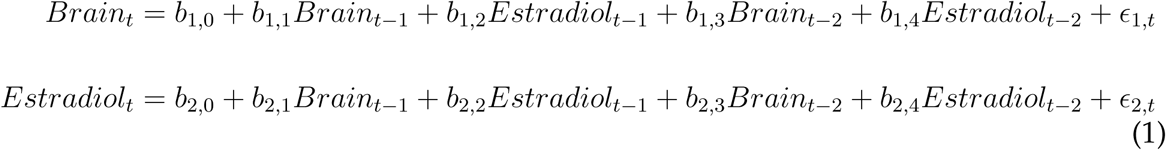

where error terms, *ϵ*_*i,t*_, are assumed to be uncorrelated and normally-distributed. Given that the design matrix is identical for each outcome measure, they can be combined in matrix form, and a least-squares solution to the system of equations can be obtained via maximum likelihood.

With respect to brain states, we modeled both edgewise coherence and factors related to macroscale network topologies. Specifically, we computed measures of *between-network* integration (the participation coefficient; i.e. the average extent to which network nodes are communicating with other networks over time) and *within-network* integration (global efficiency, quantifying the ostensible ease of information transfer across nodes inside a given network). These were derived using the relevant functions for weighted graphs in the Brain Connectivity toolbox (Rubinov and Sporns, 2010). Estimation of participation coefficients took the full (415 × 415) FDR-thresholded coherence matrices along with a vector of network IDs, quantifying the extent to which each node was connected to other nodes outside of its own network; summary, mean participation coefficients were then obtained for each network across its constituent nodes. For global efficiencies, the 415 × 415 matrices were subdivided into smaller network-specific matrices as defined by our parcellation, yielding estimates of integration only among within-network nodes. Ultimately, regardless of brain measure, each VAR was estimated similarly to the time-synchronous analyses described above: data were *Z*-scored, models were fit, and model-level stats (test-statistics, *R*^2^, and *RMSE*) were empirically-thresholded against 10,000 iterations of nonparametric permutation testing. Here, however, both brain and hormonal data were permuted under the null hypothesis of temporal stochasticity (i.e. no autoregressive trends and no time-lagged dependencies between variables). As before, we did not apply additional corrections and offer Study 2 as an independent validation set.

Finally, for each set of edgewise models (time-synchronous and time-lagged), we attempted to disentangle both the general *direction* of hormone-related associations and whether certain networks were more or less *sensitive* to hormonal fluctuations. Toward that end, we took the thresholded statistical parametric maps for each model (where edges are test-statistics quantifying the magnitude of association between coherence and hormonal timeseries) and estimated *nodal association strengths* per graph theory’s treatment of signed, weighted networks. That is, positive and negative association strengths were computed independently for each of the 415 nodes by summing the suprathreshold positive/negative edges linked to them. We then simply assessed mean association strengths (*±* 95% confidence intervals) in each direction across the various networks in our parcellation.

Here, networks were defined by grouping the subnetworks of the 17-network Schaefer parcellation, such that (for example), the A, B, and C components of the Default Mode Network were treated as one network. We chose this due to the presence of a unique Temporal Parietal Network in the 17-network partition, which is otherwise subsumed by several other networks (Default Mode, Salience/Ventral Attention, and SomatoMotor) in the 7-network partition. The subcortical nodes of the Harvard-Oxford atlas were also treated as their own network, yielding a total of nine networks. These definitions were thus used for computation of participation coefficients and global efficiencies in network-level VAR models.

### 2.9 Brain data visualization

Statistical maps of edgewise coherence v. hormones were visualized using the Surf Ice software (https://www.nitrc.org/projects/surfice/).

## 3 Results

### 3.1 Endocrine assessments

Analysis of daily sex hormone (by liquid-chromatography mass-spectrometry; LC-MS) and gonadotropin (by chemiluminescent immunoassay) concentrations from Study 1 confirmed the expected rhythmic changes of a typical menstrual cycle, with a total cycle length of 27 days. Serum levels of estradiol and progesterone were lowest during menses (day 1–4) and peaked in late follicular (estradiol) and late luteal (progesterone) phases (**Figure 3; Table 1**). Progesterone concentrations surpassed 5 ng/mL in the luteal phase, signaling an ovulatory cycle (Leiva et al., 2015). In Study 2, the participant was placed on a pharmacological regimen (0.02 mg ethinyl-estradiol, 0.1 mg levonorgestrel) that chronically and selectively suppressed circulating progesterone, while leaving endogenous estradiol concentrations largely untouched. Estradiol dynamics in Study 2 (*M* = 66.2 pg/mL, range: 5–246 pg/mL) were highly similar to Study 1 (*M* = 82.8 pg/mL, range: 22–264 pg/mL; *t*(58) = −1.01, *p* = .32; **Figure S3**), offering us a second dataset in which to test the reliability of estradiol’s influence on intrinsic brain networks.

**Table 1.**
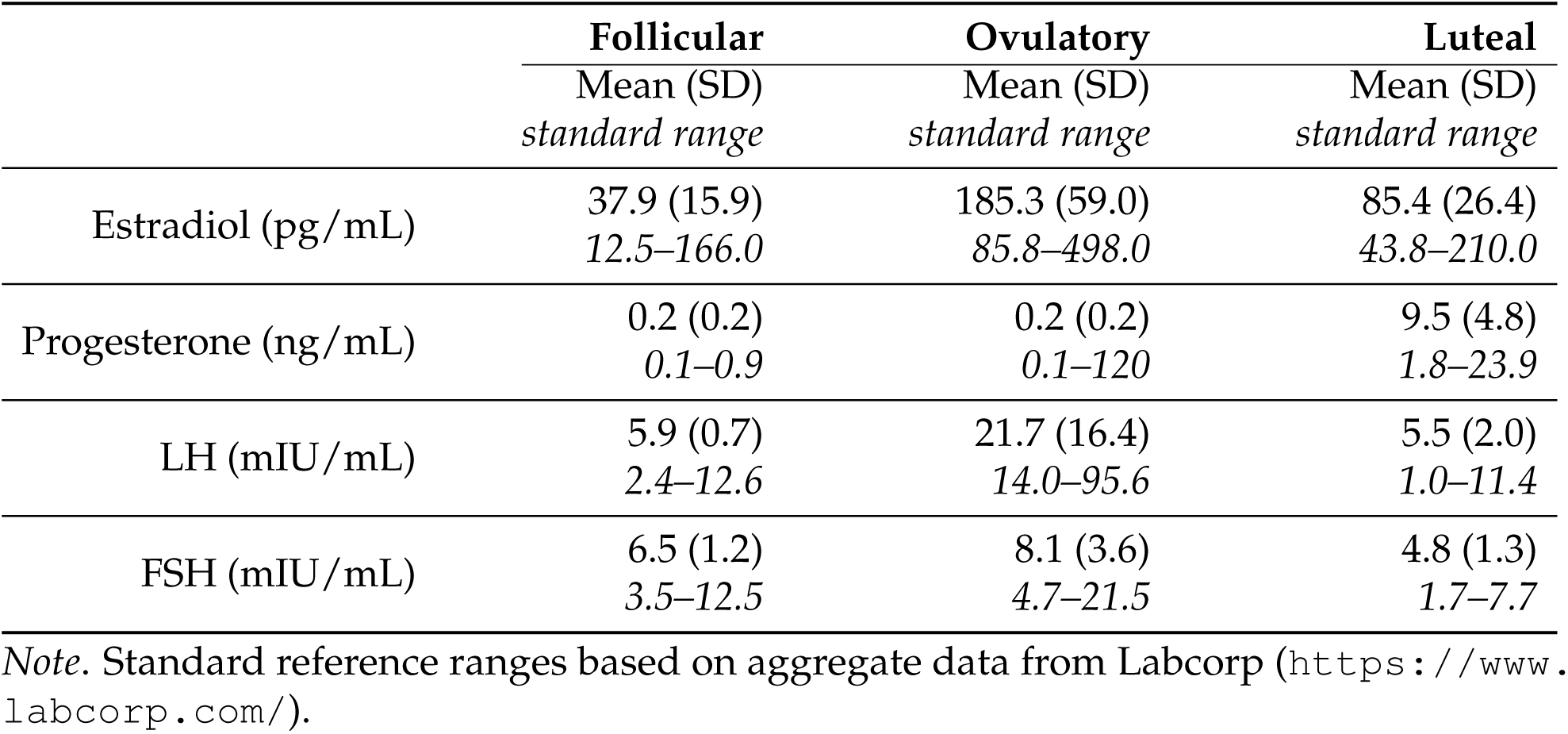
Gonadal and pituitary hormones by cycle stage (Study 1).

|  | **Follicular** | **Ovulatory** | **Luteal** |
| --- | --- | --- | --- |
|  | Mean (SD)<br>*standard range* | Mean (SD)<br>*standard range* | Mean (SD)<br>*standard range* |
| Estradiol (pg/mL) | 37.9 (15.9)<br>12.5–166.0 | 185.3 (59.0)<br>85.8–498.0 | 85.4 (26.4)<br>43.8–210.0 |
| Progesterone (ng/mL) | 0.2 (0.2)<br>0.1–0.9 | 0.2 (0.2)<br>0.1–120 | 9.5 (4.8)<br>1.8–23.9 |
| LH (mIU/mL) | 5.9 (0.7)<br>2.4–12.6 | 21.7 (16.4)<br>14.0–95.6 | 5.5 (2.0)<br>1.0–11.4 |
| FSH (mIU/mL) | 6.5 (1.2)<br>3.5–12.5 | 8.1 (3.6)<br>4.7–21.5 | 4.8 (1.3)<br>1.7–7.7 |

*Note.* Standard reference ranges based on aggregate data from Labcorp (https://www.labcorp.com/).

**Figure 3.**
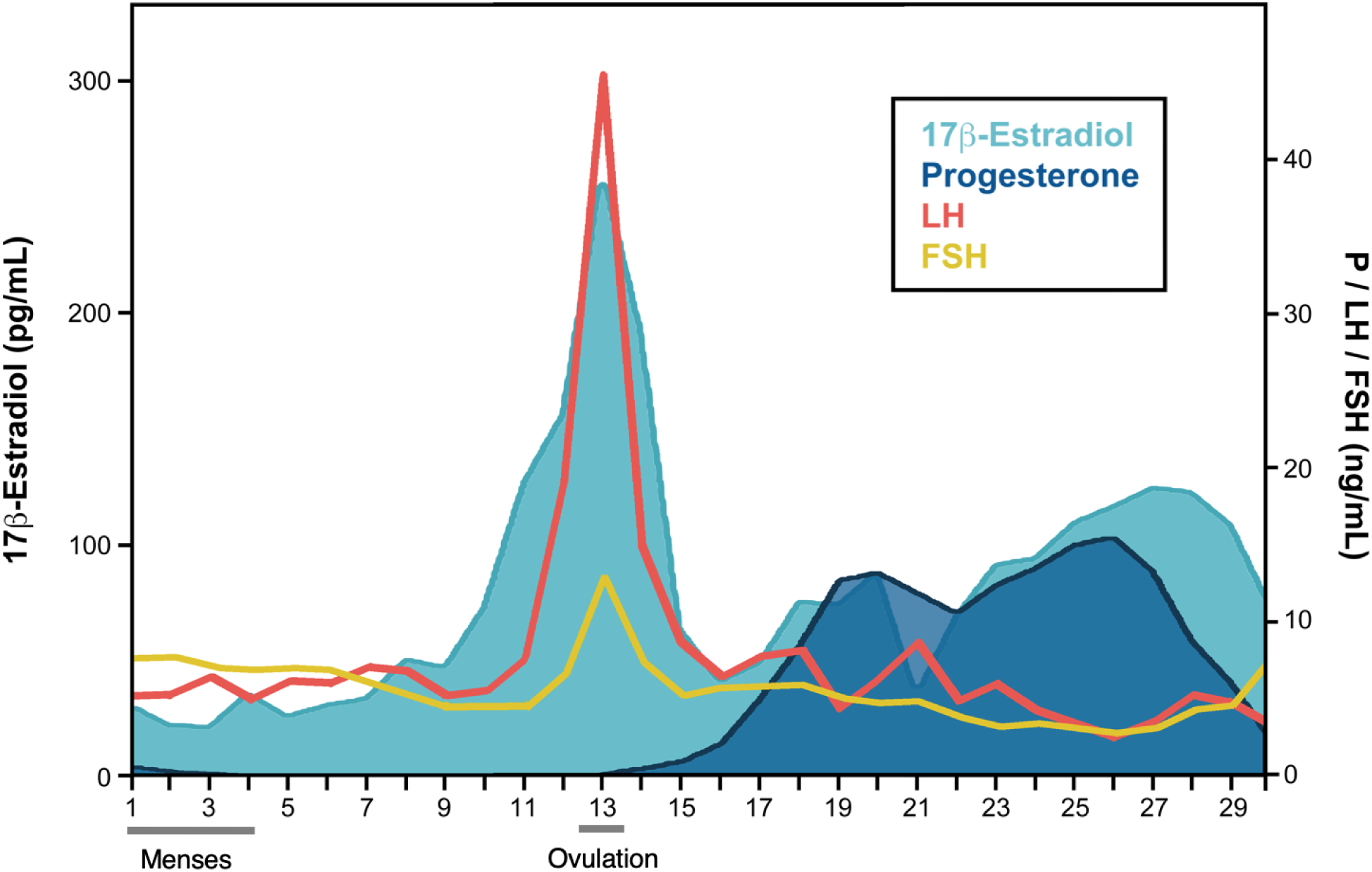
Participant’s hormone concentrations plotted by day of cycle (Study 1). 17*β*-estradiol, progesterone, luteinizing hormone (LH), and follicle stimulating hormone (FSH) concentrations fell within standard ranges.

### 3.2 Time-synchronous associations between sex hormones and whole-brain functional connectivity

Inspection of day-to-day similarity in whole-brain patterns of coherence (via pairwise Pearson correlation) revealed moderate-to-high levels of reliability between different stages of the cycle. Notably, however, one session in Study 1 (experiment day 26) was markedly dissimilar to the other sessions. Removal of this day from the analysis below did not impact the results (**Figure S4**).

To further explore cycle-dependent variability, we tested the hypothesis that whole-brain functional connectivity at rest is associated with intrinsic fluctuations in estradiol and progesterone in a *time-synchronous* (i.e. day-by-day) fashion. Based on the enriched expression of ER in frontal cortex (Wang et al., 2010), we predicted that the Default Mode, Frontoparietal Control, and Dorsal Attention Networks would be most sensitive to hormone fluctuations across the cycle.

In Study 1, we observed robust increases in coherence as a function of increasing estradiol across the brain (**Figure 4A**). When summarizing the average magnitude of association per network (as defined by our parcellation; **Figure 4C**), components of the Temporal Parietal Network had the strongest positive associations with estradiol on average, as well as the most variance (**Figure 4D**). With the exception of Subcortical nodes, all networks demonstrated some level of significantly positive association on average (95% CIs not intersecting zero). We observed a paucity of edges showing inverse associations (connectivity decreasing while estradiol increased), with no networks demonstrating significantly negative associations on average (**Figure 4D**). These findings suggest that edgewise functional connectivity is primarily characterized by increased coupling as estradiol rises over the course of the cycle.

**Figure 4.**
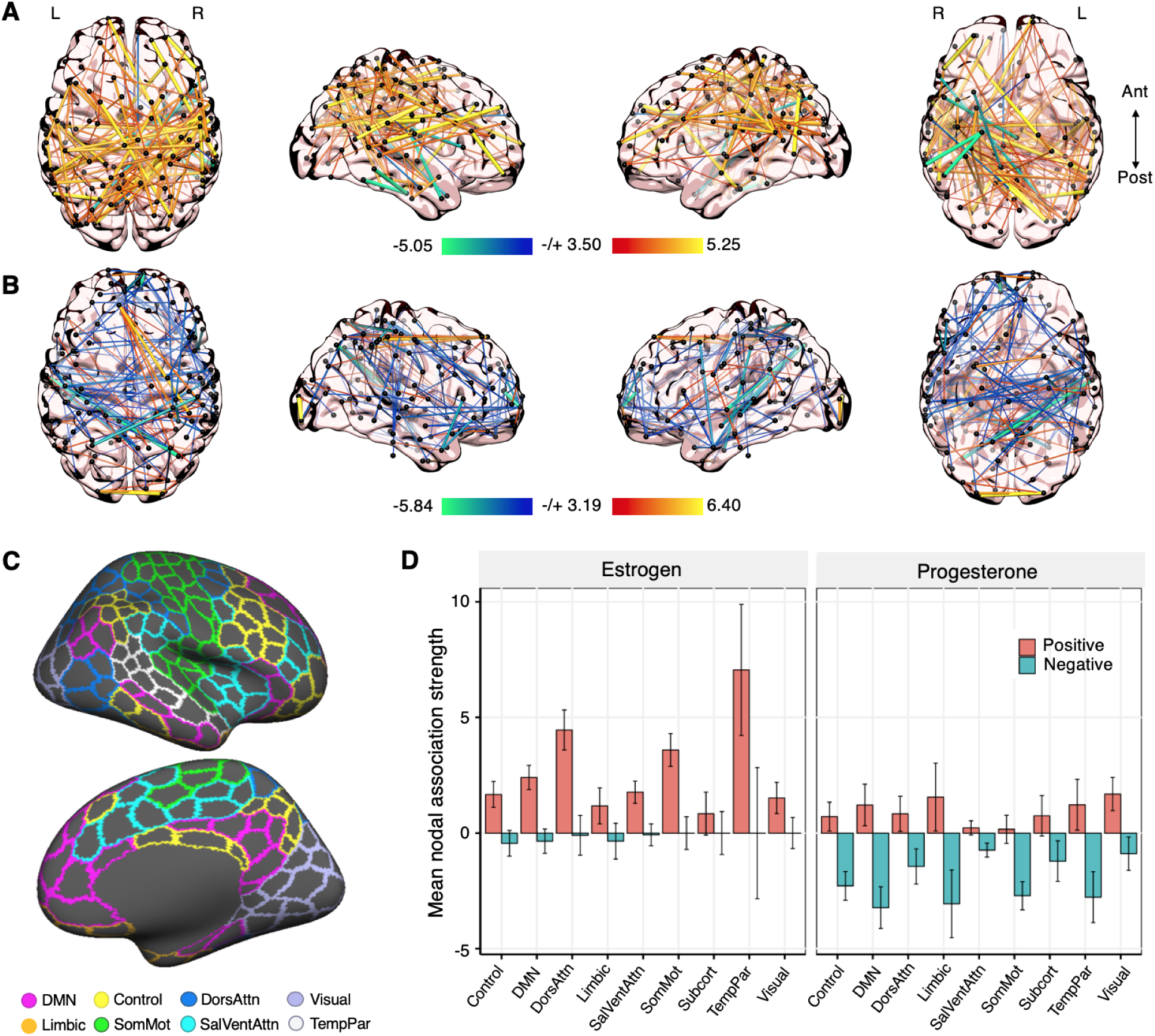
Whole-brain functional connectivity at rest is associated with intrinsic fluctuations in estradiol and progesterone (Study 1). (**A**) Time-synchronous (i.e. day-by-day) associations between estradiol and coherence. Hotter colors indicate increased coherence with higher concentrations of estradiol; cool colors indicate the reverse. Results are empirically-thresholded via 10,000 iterations of nonparametric permutation testing (*p* < .001). Nodes without significant edges are omitted for clarity. (**B**) Time-synchronous associations between progesterone and coherence. (**C**) Cortical parcellations were defined by the 400-node Schaefer atlas (shown here). An additional 15 subcortical nodes were defined from the Harvard-Oxford atlas. (**D**) Mean nodal association strengths by network and hormone. Error bars give 95% confidence intervals. ‘Positive’ refers to the average magnitude of positive associations (e.g. stronger coherence with higher estradiol); ‘Negative’ refers to the average magnitude of inverse associations (e.g. weaker coherence with higher estradiol). Abbreviations: DMN, Default Mode Network; DorsAttn, Dorsal Attention Network; SalVentAttn, Salience/Ventral Attention Network; SomMot, SomatoMotor Network; TempPar, Temporal Parietal Network.

Progesterone, by contrast, yielded a widespread pattern of inverse association across the brain, such that connectivity decreased as progesterone rose (**Figure 4B**). Most networks (with the exception of the Salience/Ventral Attention and SomatoMotor Networks) still yielded some degree of significantly positive association over time; however, the general strength of negative associations was larger in magnitude and significantly nonzero across all networks (**Figure 4D**). Together, the direction of these observed relationships offers a macroscale analogue to cellular-level animal models of estradiol and progesterone function, consistent with proliferative (increased connectivity) and reductive (decreased connectivity) effects, respectively. Re-analysis with global signal regression included during preprocessing yielded a similar pattern of results (**Figure S5**), suggesting that the relationships observed in Study 1 are *not* due to arbitrary changes in global signal over time (e.g. due to physiological variability over the cycle).

Given the predominantly positive associations between connectivity and estradiol, we further assessed the dependence of these effects on the estradiol surge that occurs during ovulation. Removal of the ovulation window erased significant associations across the brain almost entirely (**Figure S6A**), indicating that the hallmark rise of estradiol during ovulation may be a key modulator of functional coupling over a reproductive cycle. We then tested the reliability of these associations when estradiol fluctuations were unopposed by progesterone (Study 2): this revealed similarly ubiquitous increases in connectivity coincident with estradiol fluctuations (**Figure S7**). As before, removal of the three highest estradiol days during the mid-cycle peak (akin to the ovulatory window from Study 1) greatly reduced whole-brain associations (**Figure S6B**). Thus, whole-brain functional connectivity appears highly-sensitive to estradiol regardless of reproductive status.

### 3.3 Time-lagged associations between estradiol and whole-brain functional connectivity

We then employed time-lagged methods from dynamical systems analysis to further elucidate the degree to which intrinsic functional connectivity is sensitive to fluctuations in estradiol: specifically, vector autoregression (VAR), which supports more *directed* temporal inference than standard regression models. As described previously, we report results from second-order VAR models: thus, in order to assess connectivity or hormonal states on a given day of the experiment, we consider their values on both the previous day (hereafter referred to as ‘lag 1’) and two days prior (hereafter referred to as ‘lag 2’). Ultimately, if brain variance over time is attributable to previous states of estradiol, this suggests that temporal dynamics in connectivity may be *driven* (in part) by fluctuations in this hormone.

When assessing edgewise connectivity states, a powerful disparity emerged between the brain’s autoregressive effects and the effects of estradiol in Study 1. We observed vast, whole-brain associations with prior hormonal states, both at lag 1 and lag 2 (**Figure 5A**). Perhaps most immediately striking, the sign of these brain-hormone associations inverts between lags, such that it is predominantly positive at lag 1 and predominantly negative at lag 2—this holds for all networks when considering their mean nodal association strengths (**Figure 5B**). We interpret this as a potential regulatory dance between brain states and hormones over the course of the cycle, with estradiol perhaps playing a role in maintaining both steady states (when estradiol is low) and transiently-high dynamics (when estradiol rises). No such pattern emerged in the brain’s autoregressive effects, with sparse, low-magnitude, and predominantly negative associations at lag 1 and lag 2 (**Figure S8**). The observed associations between estradiol and edgewise connectivity were partially unidirectional. Previous states of coherence were associated with estradiol across a number of edges, intersecting all brain networks. This emerged at both lag 1 and lag 2; however, unlike the lagged effects of estradiol on coherence, association strengths were predominantly negative and low-magnitude (on average) at both lags (**Figure S9**). Moreover—and importantly—none of the edges that informed the temporal evolution of estradiol were also significantly preceded *by* estradiol at either lag (i.e. there was no evidence of mutual modulation at any network edge).

**Figure 5.**
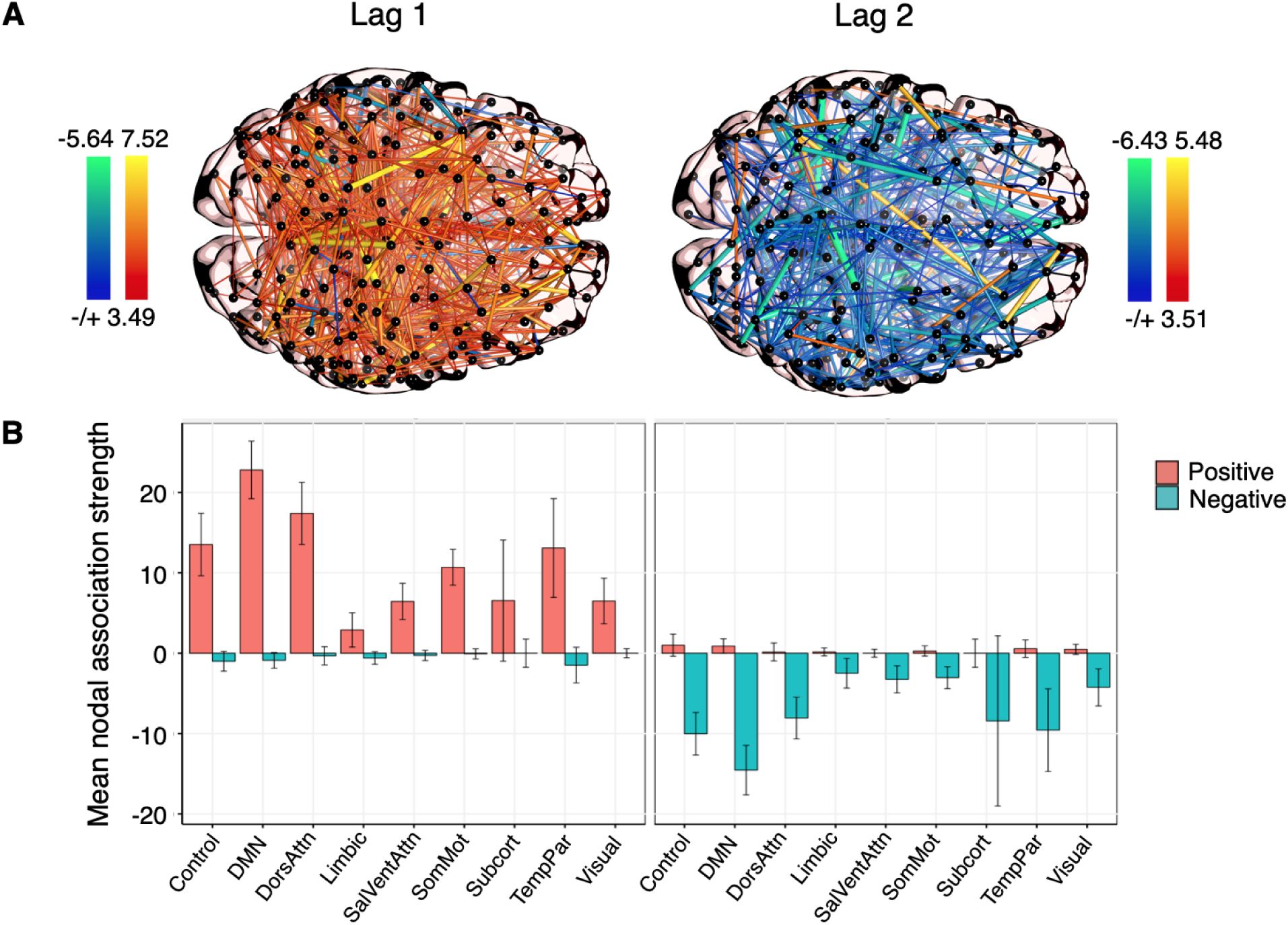
Whole-brain functional connectivity is linearly dependent on previous states of estradiol (Study 1). (**A**) Time-lagged associations between coherence and estradiol at lag 1 (*left*) and lag 2 (*right*), derived from edgewise vector autoregression models. Hotter colors indicate a predicted increase in coherence given previous concentrations of estradiol; cool colors indicate the reverse. Results are empirically-thresholded via 10,000 iterations of nonparametric permutation testing (*p* < .001). Nodes without significant edges are omitted for clarity. (**B**) Mean nodal association strengths by network and time lag. Error bars give 95% confidence intervals. ‘Positive’ refers to the average magnitude of positive associations (stronger coherence when prior states of estradiol where high); ‘Negative’ refers to the average magnitude of inverse associations (weaker coherence when prior states of estradiol were high).

We again tested the reliability of these effects in the replication sample. The autoregressive trends in edgewise coherence remained sparse and low-magnitude on average; however, unlike the original sample, nearly all networks demonstrated significantly positive associations at lag 1, and lag 2 was dominated by negative associations (**Figure S10**). Previous states of coherence also informed changes in estradiol over time, but this, too, differed from the original sample at the network level. While coherence at lag 1 was generally associated with decreases in estradiol across most networks, several networks (including the Control, Default Mode, and Dorsal Attention Networks) were associated with increases on average at lag 2 (**Figure S11**). Finally, and importantly, we observed highly-robust associations between lagged states of estradiol and coherence, with widespread positive associations at lag 1 and predominantly negative associations at lag 2 (**Figure S12**). Curiously, in contrast to the naturally-cycling data, ‘non-cognitive’ networks such as the SomatoMotor and Visual Networks demonstrated by far the strongest-magnitude associations on average—particularly at lag 1. It is possible that estradiol’s effects are magnified when unopposed by the inhibitory nature of progesterone, a topic to be addressed in future investigations.

### 3.4 Time-lagged associations between estradiol and functional network topologies

Given the findings above, we applied the same time-lagged framework to *topological states* of brain networks in order to better capture the directionality and extent of brain-hormone interactions at the mesoscopic level. These states were quantified using common graph theory metrics: namely, the *participation coefficient* (an estimate of *between-network* integration) and *global efficiency* (an estimate of *within-network* integration). We focus on significant network-level effects below, but a full documentation of our findings is available in the **Supplementary Tables**.

#### 3.4.1 Estradiol and between-network participation

As expected, estradiol demonstrated significant autoregressive trends across all models in Study 1. However, between-network integration was only tenuously associated with previous states of estradiol: in several intrinsic networks, overall model fit (variance accounted for, *R*^2^, and root mean-squared error, *RMSE*) was at best marginal compared to empirical null distributions of these statistics, and we therefore urge caution in interpreting these results. For example, in the Dorsal Attention Network (DAN; **Figure 6A-B; Table 2**), estradiol was significantly associated with between-network participation both at lag 1 (*b* = −0.56, *SE* = 0.25, *t* = −2.27, *p* = .035) and at lag 2 (*b* = 0.53, *SE* = 0.24, *t* = 2.16, *p* = .042). Overall fit for DAN participation, however, rested at the classical frequentist threshold for significance, relative to empirical nulls (*R*^2^ = 0.32, *p* = .049; *RMSE* = 0.79, *p* = .050). We observed a similar pattern of results for the Default Mode Network (DMN) and Limbic Network, where lagged states of estradiol were significantly associated with cross-network participation, but model fit as a whole was low (see **Table S1**).

**Table 2.**
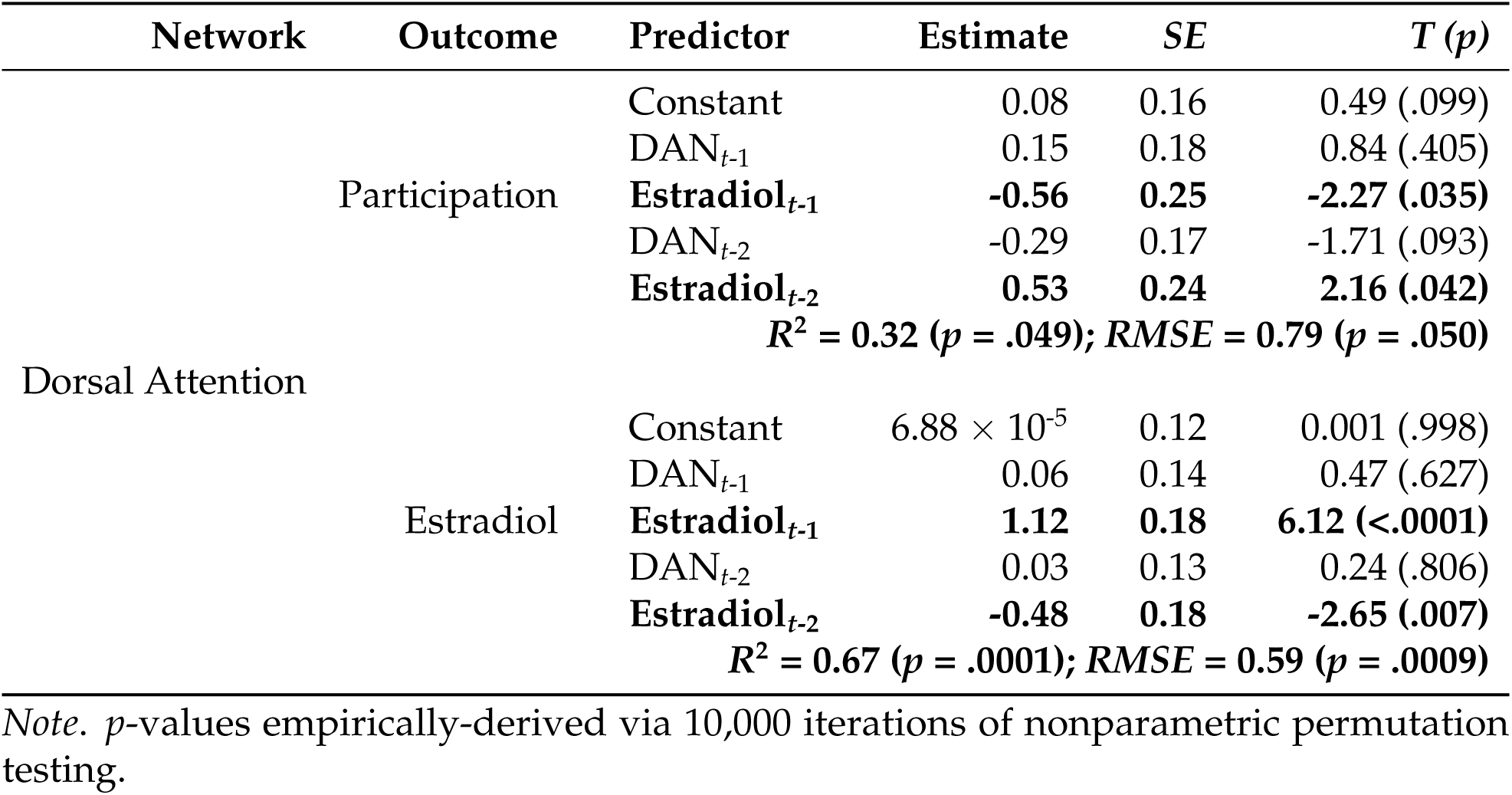
VAR model fit: Between-network participation (Study 1).

**Figure 6.**
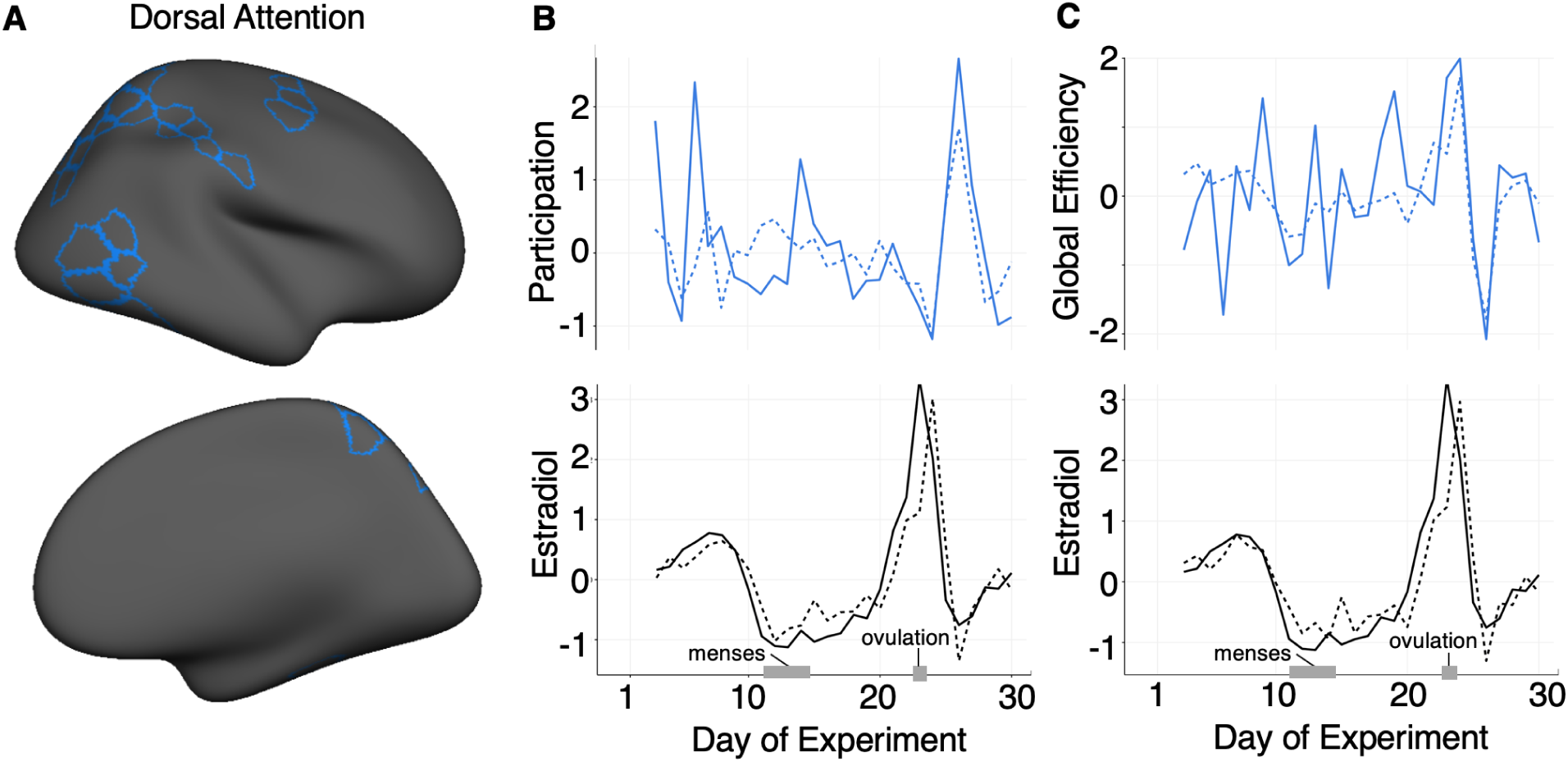
Dorsal Attention Network topology is driven by previous states of estradiol (Study 1). Observed data (solid lines) vs. VAR model fits (dotted lines) for between-network participation (**B**, *middle*) and within-network efficiency (**C**, *right*) in the Dorsal Attention Network (**A**, *left*). Timeseries for each network statistic are depicted above in (**B**,**C**) and estradiol for each VAR is plotted below. Data are in standardized units and begin at experiment day three, given the second-order VAR (lag of two days).

Importantly, we failed to replicate these effects in Study 2 under hormonal suppression (**Table S2**). The autoregressive trends in estradiol were generally blunted, with lag 2 now offering no predictive value. Previous states of DAN participation also informed the temporal evolution of estradiol (whereas estradiol predicted DAN participation in Study 1); however, this only emerged at lag 1 (*b* = −0.09, *SE* = 0.04, *t* = −2.08, *p* = .044). The Limbic and Subcortical Networks additionally demonstrated significant autoregressive trends at lag 1, but neither showed significant associations with estradiol. In sum, the marginal model fits, along with failures to replicate in Study 2, requires future investigation before robust conclusions can be drawn for between-network participation.

#### 3.4.2 Estradiol and global efficiency

In contrast to between-network integration, estradiol was more strongly associated with within-network integration, both in terms of parameter estimates and overall fit. Here, the Default Mode Network provided the best-fitting model in Study 1 (*R*^2^ = 0.50, *p* = .003; *RMSE* = 0.70, *p* = .022; **Figure 7A-B**). As before, estradiol demonstrated significant autoregressive effects at lag 1 (*b* = 1.15, *SE* = 0.19, *t* = 6.15, *p* < .0001) and lag 2 (*b* = −0.48, *SE* = 0.19, *t* = −2.50, *p* = .012). When assessing dynamics in DMN efficiency, previous states of estradiol remained significant both at lag 1 (*b* = 0.98, *SE* = 0.23, *t* = 3.37, *p* = .0003) and at lag 2 (*b* = −0.93, *SE* = 0.23, *t* = −4.00, *p* = .002). Critically, these effects were purely directional: prior states of Default Mode efficiency were not associated with estradiol, nor did they have significant autoregressive effects, suggesting that variance in topological network states (perhaps within-network integration, in particular) is primarily accounted for by estradiol—not the other way around (**Table 3**).

**Table 3.**
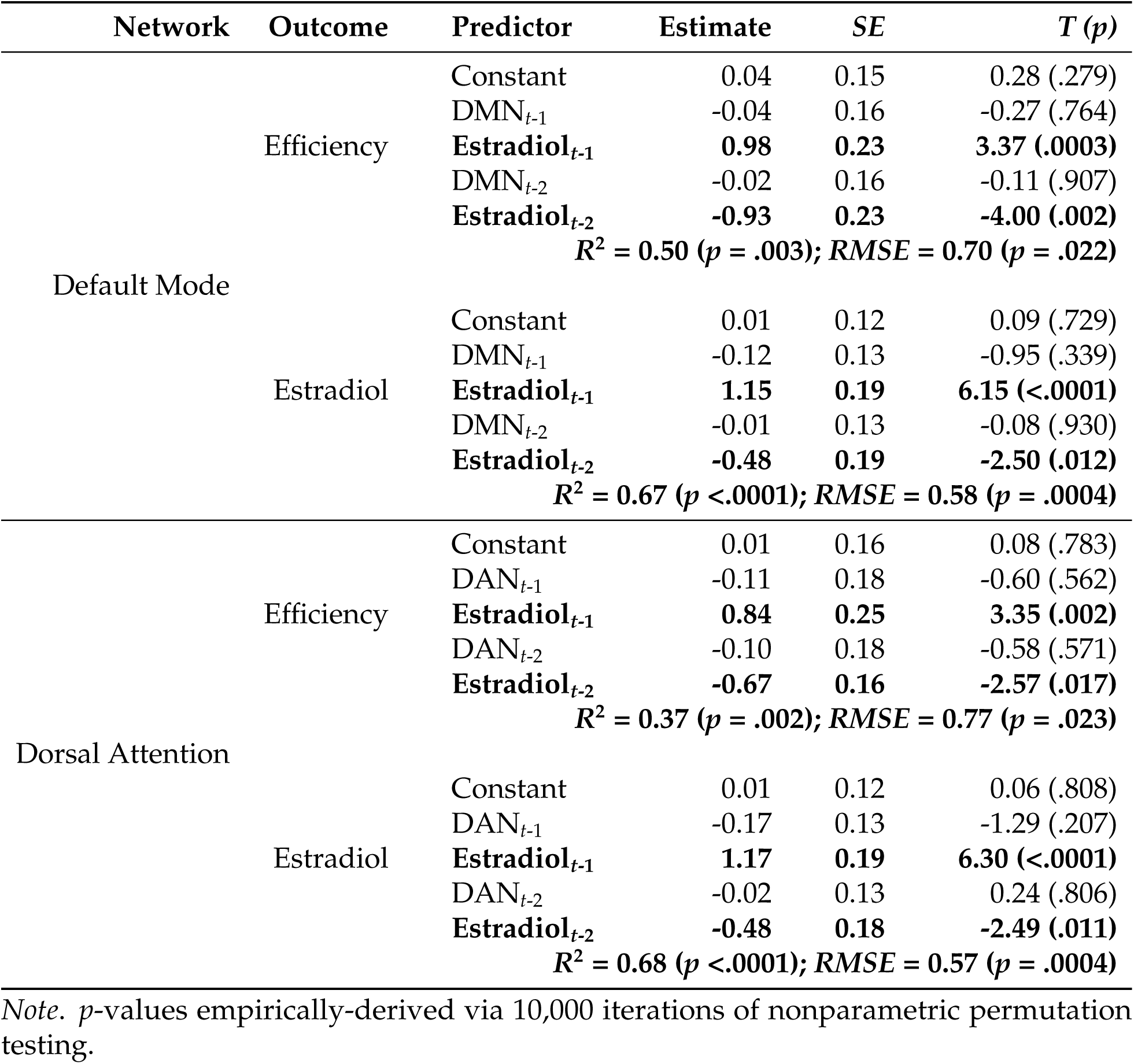
VAR model fit: Global efficiency (Study 1).

**Figure 7.**
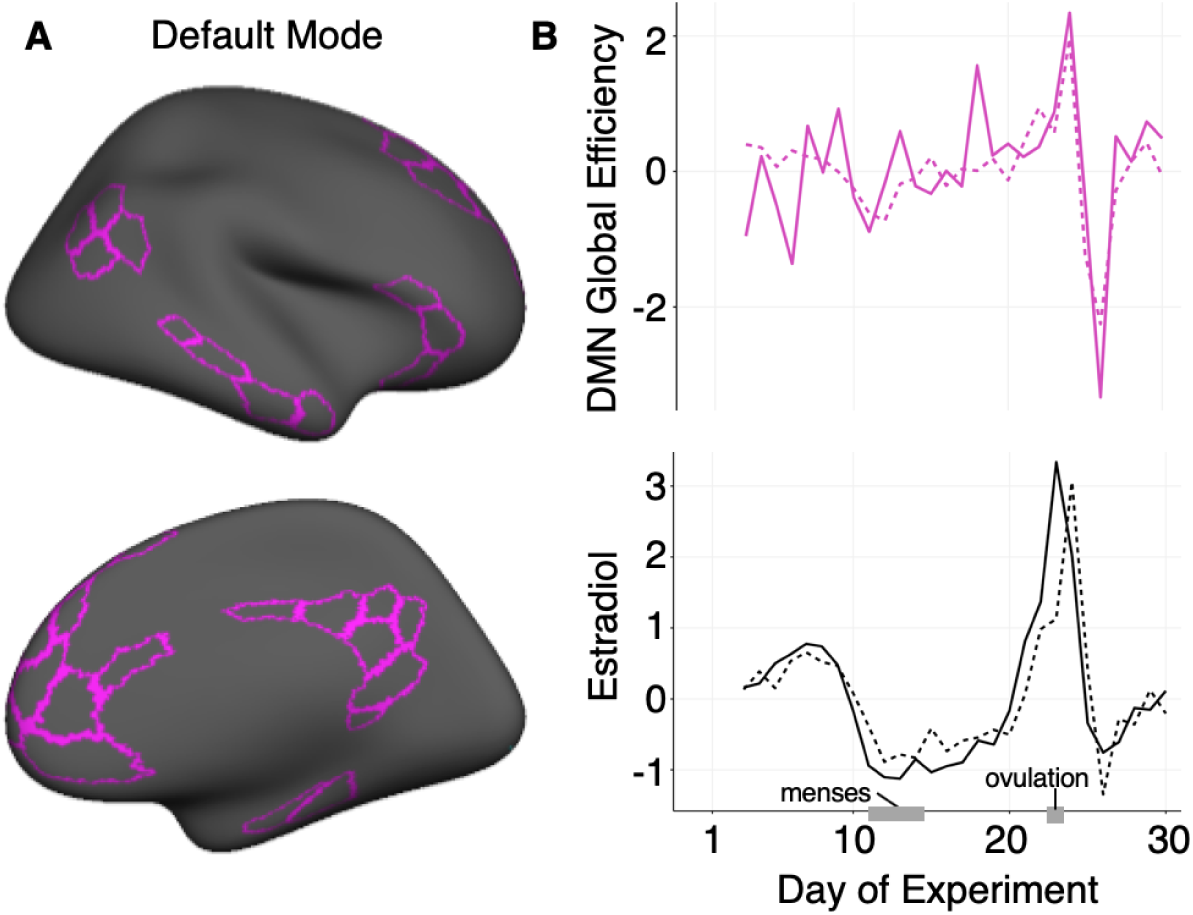
Default Mode Network topology is driven by previous states of estradiol (Study 1). Observed data (solid lines) vs. VAR model fits (dotted lines) for within-network efficiency (**B**, *right*) in the Default Mode Network (**A**, *left*). The efficiency timeseries is depicted above in (**B**) and estradiol is plotted below. Data are in standardized units and begin at experiment day three, given the second-order VAR (lag of two days).

We observed a similar pattern of results in the Dorsal Attention Network (*R*^2^ = 0.37, *p* = .022; *RMSE* = 0.77, *p* = .023; **Figure 6C; Table 3**). Estradiol again demonstrated significant autoregressive trends at lag 1 (*b* = 1.17, *SE* = 0.19, *t* = 6.30, *p* < .0001) and lag 2 (*b* = −0.48, *SE* = 0.19, *t* = −2.49, *p* = .011), as well as significant lagged associations with DAN efficiency both at lag 1 (*b* = 0.84, *SE* = 0.25, *t* = 3.35, *p* = .002) and at lag 2 (*b* = −0.67, *SE* = 0.16, *t* = −2.57, *p* = .017). As above, Dorsal Attention efficiency had no significant effects on estradiol, nor were there significant autoregressive effects of the network on itself.

The Control and Temporal Parietal networks also yielded partial support for time-dependent modulation of efficiency by estradiol (Control *R*^2^ = 0.34, *p* = .039; Temporal Parietal *R*^2^ = 0.36, *p* = .026). The time-lagged effects of estradiol followed the trends observed above; however, the overall model fit (with respect to prediction error) was not significantly better than their empirical nulls (Control *RMSE* = 0.83, *p* = .133; Temporal Parietal *RMSE* = 0.79, *p* = .057). Estradiol did not explain a significant proportion of variance in efficiency for any other networks in Study 1 (**Table S3**).

In contrast to between-network participation, within-network efficiency yielded stronger evidence for replication in Study 2. The DMN again demonstrated the strongest model fit (*R*^2^ = 0.38, *p* = .019; *RMSE* = 0.74, *p* = .011), with estradiol informing fluctuations in DMN efficiency both at lag 1 (*b* = 2.48, *SE* = 0.75, *t* = 3.29, *p* = .003) and lag 2 (*b* = −2.69, *SE* = 0.91, *t* = −2.94, *p* = .009). We also observed a significant autoregressive effect of DMN efficiency at lag 2 (*b* = −0.45, *SE* = 0.19, *t* = −2.41, *p* = .027), but not at lag 1. In the DAN, significant model fit was achieved with respect to prediction error (*RMSE* = 0.79, *p* = .045), but variance accounted for was marginal relative to empirical nulls (*R*^2^ = 0.32, *p* = .052). Accordingly, estradiol significantly associated with DAN efficiency at lag 1 (*b* = 1.88, *SE* = 0.79, *t* = 2.37, *p* = .026) but not at lag 2. Finally, previous states of estradiol (both lags 1 and 2) significantly informed efficiency in the Control, Salience / Ventral Attention, SomatoMotor, and Subcortical Networks; however, aside from the SomatoMotor Network (*R*^2^ = 0.34, *p* = .039; *RMSE* = 0.76, *p* = .018), overall fit in these models was nonsignificant (**Table S4**). Thus, while we observed trends largely consistent with Study 1 (with respect to DMN and DAN efficiency), there may be additional network-level effects in a neuroendocrine system unopposed by progesterone, warranting future investigation.

## 4 Discussion

In this dense-sampling, deep-phenotyping project, a naturally-cycling female underwent resting state fMRI and venipuncture for 30 consecutive days, capturing the dynamic endocrine changes that unfold over the course of a complete menstrual cycle. Time-synchronous analyses illustrate estradiol’s widespread associations with cortical network dynamics, spanning all but one of the networks in our parcellation. Time-lagged vector autoregressive models tested the temporal directionality of these effects, suggesting that intrinsic network dynamics may be partially driven by recent states of estradiol, particularly with respect to within-network connectivity: global efficiency in the Default Mode and Dorsal Attention Networks exhibited the strongest associations with fluctuations in estradiol, replicated between Studies 1 and 2. In contrast to estradiol’s proliferative effects, progesterone was primarily associated with reduced coherence across the whole brain. Findings from a replication dataset further establish estradiol’s impact on large-scale cortical dynamics. Critically, removal of high estradiol days in both studies reduced associations across the brain, suggesting that the hallmark rise of estradiol surrounding the ovulatory window may be a key modulator of functional coupling during the reproductive cycle (**Figure S6**). These results reveal the rhythmic nature in which brain networks reorganize across the human menstrual cycle.

The network neuroscience community has begun to probe functional networks over the timescale of weeks, months, and years to understand the extent to which brain networks vary between individuals or within an individual over time (Betzel et al., 2019; Finn et al., 2015; Gordon et al., 2017; Horien et al., 2019; Poldrack et al., 2015; Seitzman et al., 2019). These studies indicate that functional networks are dominated by common organizational principles and stable individual features, especially in frontoparietal control regions (Finn et al., 2015; Gordon et al., 2017; Gratton et al., 2018a; Horien et al., 2019). An overlooked feature of these regions is that they are populated with estrogen and progesterone receptors and are exquisitely sensitivity to major changes in sex hormone concentrations (Berman et al., 1997; Hampson and Morley, 2013; Jacobs and D’Esposito, 2011; Jacobs et al., 2016a,b; Shanmugan and Epperson, 2014). Our findings demonstrate significant effects of estradiol on functional network nodes belonging to the DMN, DAN, and FCN that overlap with ER-rich regions of the brain, including medial/dorsal PFC (Wang et al., 2010; Yeo et al., 2011). This study merges the network neuroscience and endocrinology disciplines by demonstrating that higher-order processing systems are modulated by day-to-day changes in sex hormones over the timescale of one month.

### 4.1 Sex hormones regulate brain organization across species

Animal studies offer unambiguous evidence that sex steroid hormones shape the synaptic organization of the brain, particularly in regions that support higher order cognitive functions (Frick et al., 2018, 2015; Galea et al., 2017; Hara et al., 2015; Woolley and McEwen, 1993). In rodents, estradiol increases fast-spiking interneuron excitability in deep cortical layers (Clemens et al., 2019). In nonhuman primates, whose reproductive cycle length is similar to humans, estradiol increases the number of synapses in PFC (Hara et al., 2015). Recently, this body of work has also begun to uncover the functional significance of sinusoidal *changes* in estradiol. For example, estradiol’s ability to promote PFC spinogenesis in ovariectomized animals occurs *only if* the hormone add-back regime mirrors the cyclic pattern of estradiol release typical of the macaque menstrual cycle (Hao et al., 2006; Ohm et al., 2012). Pairing estradiol with cyclic administration of progesterone blunts this increase in spine density (Ohm et al., 2012). In the hippocampus, progesterone has a similar inhibitory effect on dendritic spines, blocking the proliferative effects of estradiol 6 hours after administration (Woolley and McEwen, 1993). Together, the preclinical literature suggests that progesterone antagonizes the largely proliferative effects of estradiol (for review, see Brinton et al. (2008)). We observed a similar relationship, albeit at a different spatiotemporal resolution, with estradiol demonstrating positive associations with coherence across numerous cortical networks and progesterone having an opposite, negative association on average. In sum, animal studies have identified estradiol’s influence on regional brain organization at the microscopic scale. Here, we show that estradiol and progesterone may have analogous effects evident at the mesoscopic scale of whole-brain connectivity, measured by spectral coherence, and macroscopic features of network topology.

### 4.2 Resting-state network characteristics differ by cycle stage

Group-based and sparser-sampling neuroimaging studies provide further support that cycle stage and sex hormones impact resting state networks (De Bondt et al., 2015; Lisofsky et al., 2015; Petersen et al., 2014; Syan et al., 2017; Weis et al., 2019). For instance, Petersen et al. (2014) demonstrated that women sampled in the follicular stage had greater connectivity within default mode and execute control networks compared to those sampled in the luteal stage. Lisofsky et al. (2015) studied women four times across their menstrual cycles, observing significant increases in connectivity between the hippocampus and superior parietal lobule during the late follicular phase. However, recent work by Weis et al. (2019) provides compelling yet contrasting evidence for sex hormones’ relationships with resting-state functional connectivity: studying women three times across the cycle, their group observed heightened connectivity between a region of the left frontal cortex and the DMN during menstruation when estradiol levels are lowest. Inconsistencies between studies could be due to a number of factors such as differences in cycle staging methods, divergent functional connectivity estimation methods, or unaccounted for intra/inter-individual variability (Beltz and Moser, 2019). Our results suggest that failure to properly capture the complete ovulatory window, when estradiol levels rapidly rise, could lead to skewed estimates of stability within functional brain networks across the menstrual cycle (Hjelmervik et al., 2014). As such, dense-sampling studies provide a novel solution to capturing pivotal moments experienced across a complete human menstrual cycle. Arélin et al. (2015) sampled an individual every 2-3 days across four cycles and found that progesterone was associated with increased connectivity between the hippocampus, dorsolateral PFC and sensorimotor cortex, providing compelling evidence that inter-regional connectivity varies over the cycle. This particular dense-sampling approach allowed the authors to examine brain-hormone relationships while accounting for intra-individual cycle variation.

Estradiol is capable of inducing rapid, non-genomic effects and slower, genomic-effects on the central nervous system. For example, spine density on hippocampal neurons varies by ∼30% over the rodent estrous cycle. In-vivo MRI evidence in mice demonstrates that these hormone-mediated changes can occur rapidly, with differences detectable within a 24-hour period. To capture time-synchronous (rapid) and time-lagged (delayed) effects of sex steroid hormones, we expanded upon the approach of Arélin et al. (2015) by sampling an individual every 24 hours for 30 consecutive days. Our results illuminate how time-synchronous correlations and time-lagged computational approaches reveal unique aspects of where and how hormones exert their effect on the brain’s intrinsic networks. Time-synchronous analyses illustrated contemporaneous, zero-lag associations between estradiol, progesterone, and whole-brain connectivity. The introduction of lagged states in VAR allowed us to examine the temporal directionality of those relationships, suggest that recent fluctuations in estradiol (within two days) inform current brain states—this raises the interesting possibility that estradiol may play a partial role in driving changes in connectivity, particularly in the DMN and DAN.

### 4.3 Neurobiological interpretations of hormonal effects and future studies

The following considerations could enhance the interpretation of these data. First, our investigation deeply sampled a single woman, limiting our ability to generalize these findings to other individuals. To enrich our understanding of the relationship between sex hormones and brain function, this dense-sampling approach should be extended to a diverse sample of women. Doing so will allow us to examine the consistency of our results with respect to inter-individual differences in network organization over the menstrual cycle. Additionally, examining network organization during a state of complete hormone suppression would afford a valuable comparison, as certain oral hormonal contraceptives suppress the production of *both* ovarian hormones. If dynamic changes in estradiol are *facilitating* increases in resting connectivity, we expect hormonally-suppressed individuals to show less dynamic modulation of functional brain networks over time. Given the widespread use of oral hormonal contraceptives (100 million users worldwide), it is critical to determine whether sweeping changes to an individual’s endocrine state impacts brain states and whether this, in turn, has any bearing on cognition.

Second, in freely-cycling individuals, sex hormones function as proportionally-coupled *nonlinear* oscillators (Boker et al., 2014). Within-person cycle variability is almost as large as between-person cycle variability, which hints that there are highly-complex hormonal interactions within this regulatory system (Boker et al., 2014; Fehring et al., 2006). The VAR models we have explored reveal *linear* dependencies between brain states and hormones, but other dynamical systems methods (e.g. coupled latent differential equations) may offer more biophysical validity (Boker et al., 2014). However, the current sample size precludes robust estimation of such a model.

Third, while permutation tests have been used as empirical null models for VAR (Hyvärinen et al., 2010) and its statistical relatives (e.g. Granger causality; Barnett and Seth (2014)), the practice of temporally-scrambling a timeseries will drastically alter its autocorrelative structure and potentially skew observed dependencies over time. Phase-shifting, surrogate data tests such as the amplitude adjusted Fourier transform (AAFT) may offer more robust null distributions. However, AAFT also makes strong distributional assumptions about the original timeseries (Gaussian normality) that, unfortunately, are not met by these data. Additionally, the small sample size over a single cycle precludes the ability to derive robust surrogate realizations of the timeseries. While AAFT is arguably an *ideal* procedure for analyses such as those reported here, these data simply cannot meet the assumptions required for valid surrogate testing and thus is a major limitation within the current study. Future investigations involving larger samples of women over several cycles that allow implementation of such models will be critical.

Fourth, while coherence is theoretically robust to timing differences in the hemodynamic response function, hormones can affect the vascular system (Krause et al., 2006). Therefore, changes in coherence may be due to vascular artifacts that affect the hemodynamic response in fMRI, rather than being *neurally*-relevant. Future investigations exploring the assumptions of hemodynamics in relation to sex steroid hormone concentrations will add clarity as to how the vascular system’s response to hormones might influence large-scale brain function.

Fifth, these findings contribute to an emerging body of work on estradiol’s ability to enhance the efficiency of PFC-based cortical circuits. In cycling women performing a working memory task, PFC activity is exaggerated under low estradiol conditions and reduced under high estradiol conditions (Jacobs and D’Esposito, 2011). The same pattern is observed decades later in life: as estradiol production decreases over the menopausal transition, working memory-related PFC activity becomes more exaggerated, despite no difference in working memory performance (Jacobs et al., 2016a). Here, we show that day-to-day changes in estradiol enhance the global efficiency of functional networks, with pronounced effects in networks (DMN and FCN) that encompass major regions of the PFC (Schaefer et al., 2018; Yeo et al., 2011). Together, these findings suggest that estradiol generates a neurally efficient PFC response at rest and while engaging in a cognitive task. Estradiol’s action may occur by enhancing dopamine synthesis and release (Creutz and Kritzer, 2002). The PFC is innervated by midbrain dopaminergic neurons that form the mesocortical dopamine track (Kritzer and Creutz, 2008). Decades of evidence have established that dopamine signaling enhances the signal-to-noise ratio of PFC pyramidal neurons (Williams and Goldman-Rakic, 1995) and drives cortical efficiency (Cai and Arnsten, 1997; Gibbs and D’Esposito, 2005; Granon et al., 2000; Vijayraghavan et al., 2007). In turn, estradiol enhances dopamine release and modifies the basal firing rate of dopaminergic neurons (Becker, 1990; Pasqualini et al., 1995; Thompson and Moss, 1994), a possible neurobiological mechanism by which alterations in estradiol could impact cortical efficiency. Future multimodal neuroimaging studies in humans can clarify the link between estradiol’s ability to stimulate dopamine release and the hormone’s ability to drive cortical efficiency within PFC circuits.

Sixth, we observed surprisingly few autoregressive effects in brain measures across our time-lagged models. This was despite relatively strong day-to-day similarity in whole-brain patterns of connectivity (**Figure S3**), and clear evidence for autocorrelation when assessing the brain data in an independent, univariate fashion. Thus, the incorporation of sex hormones into a time-lagged modeling framework attributed more temporal variability in the brain to fluctuations in hormone concentrations. Nevertheless, an ongoing debate within the network neuroscience community surrounds test-retest reliability in resting-state functional connectivity analyses. Some studies state that large amounts of data (> 20 minutes) are necessary for test-retest reliability (Gratton et al., 2018a; Noble et al., 2017), while others argue that reliability can be derived from shorter (5-15 minutes) scans (Birn et al., 2013; Van Dijk et al., 2010). We are limited in our ability to assess whether the ostensibly-weak autoregressive trends suggested by our time-lagged models would be replicated under longer scanning durations and hope future work addresses this issue.

Finally, we chose to apply a well-established group-based atlas (Schaefer et al., 2018) to improve generalizability to other individuals, as a key goal of our investigation was to demonstrate how sex steroid hormones explain variability in intrinsic network topologies based on regional definitions shown to be reliable across thousands of individuals (Schaefer et al., 2018; Yeo et al., 2011). Yet, group-based atlases can lead to potential loss in individual-level specificity, and recent work has demonstrated that fixed atlases may not capture underlying reconfigurations in the parcellations themselves within an individual (Bijsterbosch et al., 2019; Salehi et al., 2020a,b). Therefore, future work using individual-derived functional networks will be necessary to determine whether spatial reconfigurations in *parcellations* emerge as a function of the menstrual cycle, over and above the influence of state or trait features. Relatedly, variation in analytic pipelines of brain imaging data can lead to divergent conclusions even within the same dataset (Botvinik-Nezer et al., 2020); for complete transparency, we are committed to making all neuroimaging data and code publicly available so that other investigators can assess these brain-hormone associations using their preferred methods (see **Data availability**).

### 4.4 Estradiol modulates global efficiency in estrogen-receptor rich brain regions

Using dense-sampling approaches to probe brain-hormone interactions could reveal organizational principles of the functional connectome previously unknown, transforming our understanding of how hormones influence brain states. Human studies implicate sex steroids in the regulation of brain structure and function, particularly within ER-rich regions like the PFC and hippocampus (Berman et al., 1997; Girard et al., 2017; Hampson and Morley, 2013; Jacobs and D’Esposito, 2011; Jacobs et al., 2015, 2016a,b; Shanmugan and Epperson, 2014; Zeydan et al., 2019), and yet, the neuroendocrine basis of the brain’s network organization remains understudied. Here, we used a network neuroscience approach to investigate how hormones modulate the topological integration of functional networks across the brain, showing that estradiol is associated with increased coherence across broad swaths of cortex that extend beyond regions with established ER expression. At the network level, estradiol enhances within-network efficiency (with robust effects in DAN and DMN) and, to a lesser extent, modulates between-network participation (although critically, this finding failed to replicate in Study 2). Moving forward, a complete mapping of ER/PR expression in the human brain will be essential for our understanding and interpretation of brain-hormone interactions. Furthermore, this dense-sampling approach could be applied to brain imaging studies of other major neuroendocrine transitions, such as pubertal development and reproductive aging (e.g. menopause).

### 4.5 Implications of hormonally regulated network dynamics for cognition

An overarching goal of network neuroscience is to understand how coordinated activity within and between functional brain networks supports cognition. Increased global efficiency is thought to optimize a cognitive workspace (Bullmore and Bassett, 2011), while between-network connectivity may be integral for integrating top-down signals from multiple higher-order control hubs (Gratton et al., 2018b). The dynamic reconfiguration of functional brain networks is implicated in performance across cognitive domains, including motor learning (Bassett et al., 2011; Mattar et al., 2018), cognitive control (Seeley et al., 2007), and memory (Fornito et al., 2012). Our results suggest that the within-network connectivity of these large-scale networks is temporally-dependent on hormone fluctuations across the human menstrual cycle, particularly in states of high estradiol (e.g. ovulation). Future studies should consider whether these network changes confer advantages to domain-general or domain-specific cognitive performance. Accordingly, future planned analyses from this dataset will incorporate task-based fMRI to determine whether the brain’s network architecture is similarly-variable across the cycle when engaging in a cognitive task, or in the dynamic reconfiguration that occurs when transitioning from rest to task.

### 4.6 Implications of hormonally regulated network dynamics for clinical diagnoses

Clinical network neuroscience seeks to understand how large-scale brain networks differ between healthy and patient populations (Fox and Greicius, 2010; Hallquist and Hillary, 2019). Disruptions in functional brain networks are implicated in a number of neurodegenerative and neuropsychiatric disorders: intrinsic connectivity abnormalities in the DMN are evident in major depressive disorder (Greicius et al., 2007) and Alzheimer’s disease (Buckner et al., 2009). Notably, these conditions have a sex-skewed disease prevalence: women are at twice the risk for depression and make up two-thirds of the Alzheimer’s disease patient population (Nebel et al., 2018). Here, we show that estradiol modulates efficiency within the DMN and DAN, with pronounced rises in estradiol significantly preceding increases in within-network coherence. A long history of clinical evidence further implicates sex hormones in the development of mood disorders (Plotsky et al., 1998; Rubinow and Schmidt, 2006; Young and Korszun, 2002). For example, the incidence of major depression increases with pubertal onset in females (Angold and Costello, 2006), chronic use of hormonal contraceptives (Young et al., 2007), the postpartum period (Bloch et al., 2000), and perimenopause (Schmidt and Rubinow, 2009). Moving forward, a network neuroscience approach might have greater success at identifying the large-scale network disturbances that underlie, or predict, the emergence of disease symptomology by incorporating sex-dependent variables (such as endocrine status) into clinical models. This may be particularly true during periods of profound neuroendocrine change (e.g. puberty, pregnancy, menopause, and use of hormone-based medications, reviewed by Taylor et al. (2019)) given that these hormonal transitions are associated with a heightened risk for mood disorders.

## 5 Conclusion

In sum, endogenous hormone fluctuations over the reproductive cycle have a robust impact on the intrinsic network properties of the human brain. Despite over 20 years of evidence from rodent, nonhuman primate, and human studies demonstrating the tightly-coupled relationship between our endocrine and nervous systems (Beltz and Moser, 2019; Hara et al., 2015; McEwen, 2018), the field of network neuroscience has largely overlooked how endocrine factors shape the brain. The dynamic endocrine changes that unfold over the menstrual cycle are a natural feature of half of the world’s population. Understanding how these changes in sex hormones might influence the large-scale functional architecture of the human brain is imperative for our basic understanding of the brain and for women’s health.

## End Notes

## Acknowledgements

This work was supported by the Brain and Behavior Research Foundation (EGJ), the California Nanosystems Institute (EGJ), the Institute for Collaborative Biotechnologies through grant W911NF-19-D-0001 from the U.S. Army Research Office (MBM), and the Rutherford B. Fett Fund (STG). Thanks to Mario Mendoza for phlebotomy and MRI assistance. We would also like to thank Courtney Kenyon, Maggie Hayes, and Morgan Fitzgerald for assistance with data collection.

## Author contributions

The overall study was conceived by L.P., C.M.T., S.T.G., and E.G.J.; L.P., T.S., E.L., C.M.T., S.Y., and E.G.J. performed the experiments; data analysis strategy was conceived by T.S. and L.P. and implemented by T.S.; L.P., T.S., and E.G.J. wrote the manuscript; E.L., C.M.T., S.Y., M.B.M., and S.T.G. edited the manuscript.

## Data/code availability

The full dataset consists of daily mood, diet, and behavioral assessments, task-based and resting-state fMRI, structural MRI, and serum assessments of pituitary gonadotropins and ovarian sex hormones. The dataset and all analysis code will be made publicly available upon publication.

