## Supplementary Information for "Functional reorganization of brain networks across the human menstrual cycle"

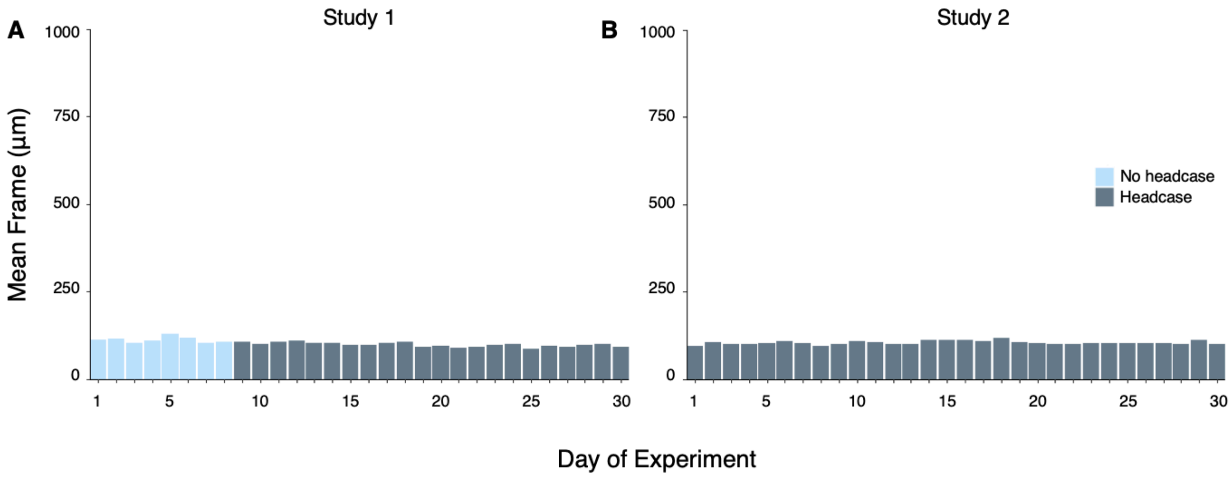


**Figure S1. Head motion (mean framewise displacement) across each 30-day experiment.** In Study 1 (**A**), motion on days 1-8 was limited using ample head and neck padding; on days 9-30, motion was limited using a molded headcase custom-fit to the participant's head. For both Study 1 and 2, mean framewise displacement did not exceed 130 microns.

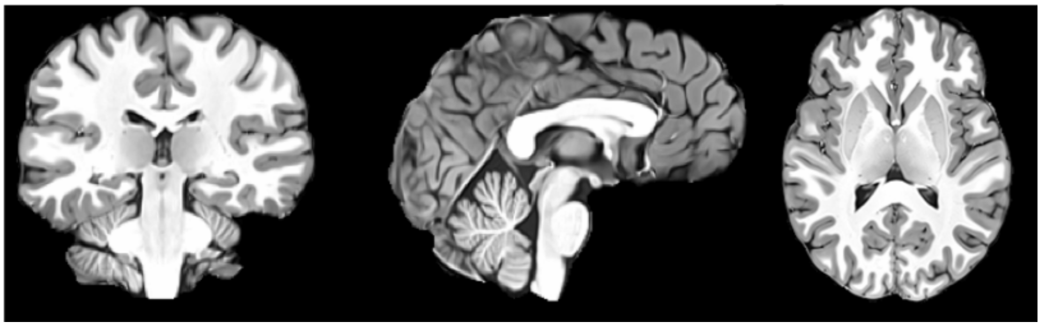

**Figure S2. Subject-specific anatomical template.** Functional images were registered to a subject-specific anatomical template space, created with Advanced Normalization Tools' multivariate template construction. All 30 high-resolution $T_1$ MPRAGE scans from Study 1 were used.

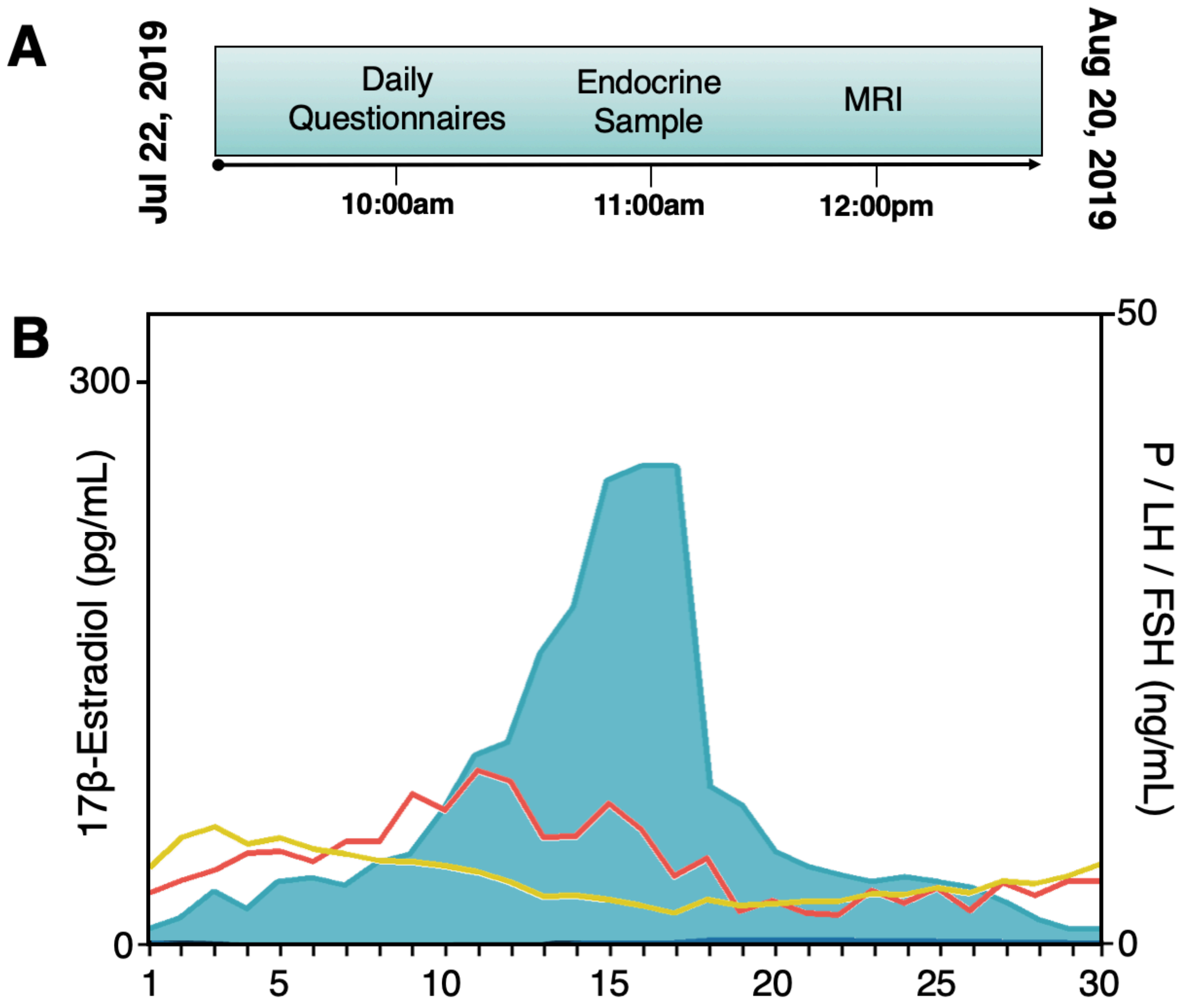


**Figure S3. Experiment timeline and hormone concentrations in Study 2. (A)** Endocrine and MRI data acquisition were again time-locked each day for 30 consecutive days. **(B)** Levels of estradiol (light blue), LH (red) and FSH (yellow) over the course of the experiment. Note that, while progesterone (dark blue) was pharmacologically suppressed, estradiol maintained a similar pattern of fluctuation as seen in Study 1.

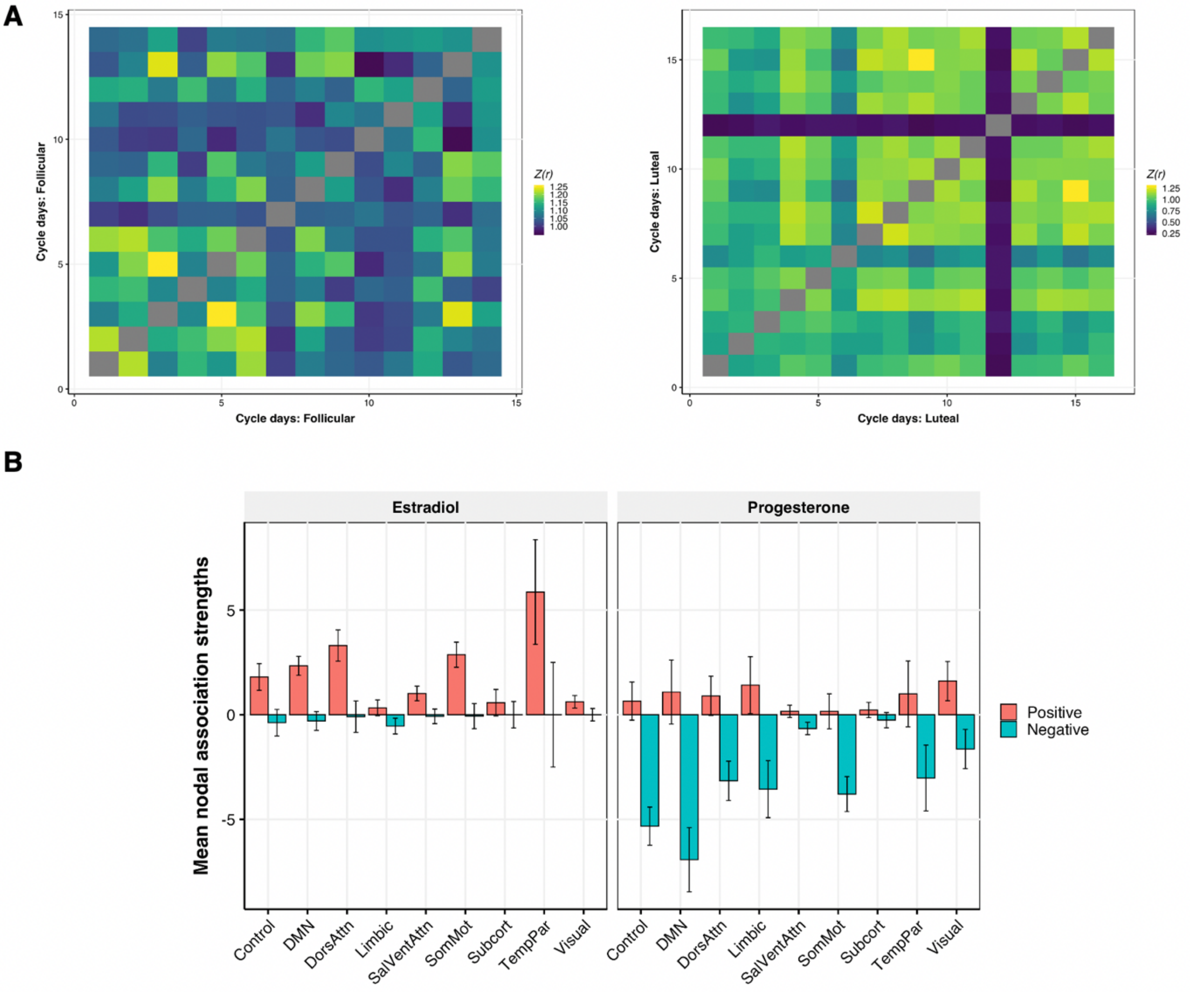


**Figure S4. Whole-brain network similarity by stage of menstrual cycle. (A)** Experiment days were divided into follicular (*left*) and luteal (*right*) stages of the cycle, and unthresholded coherence matrices were pairwise correlated with each other in order to assess inter-scan similarity in whole-brain patterns of connectivity. Fisher-*Z* transformed estimates are given. This revealed one session (experiment day 26) in the luteal stage that was markedly dissimilar relative to other days. **(B)** Mean nodal association strengths by network and hormone, after removing day 26 from the analysis shown in **Figure 4**. Critically, results were robust to this exclusion, suggesting no strong dependence on this session.

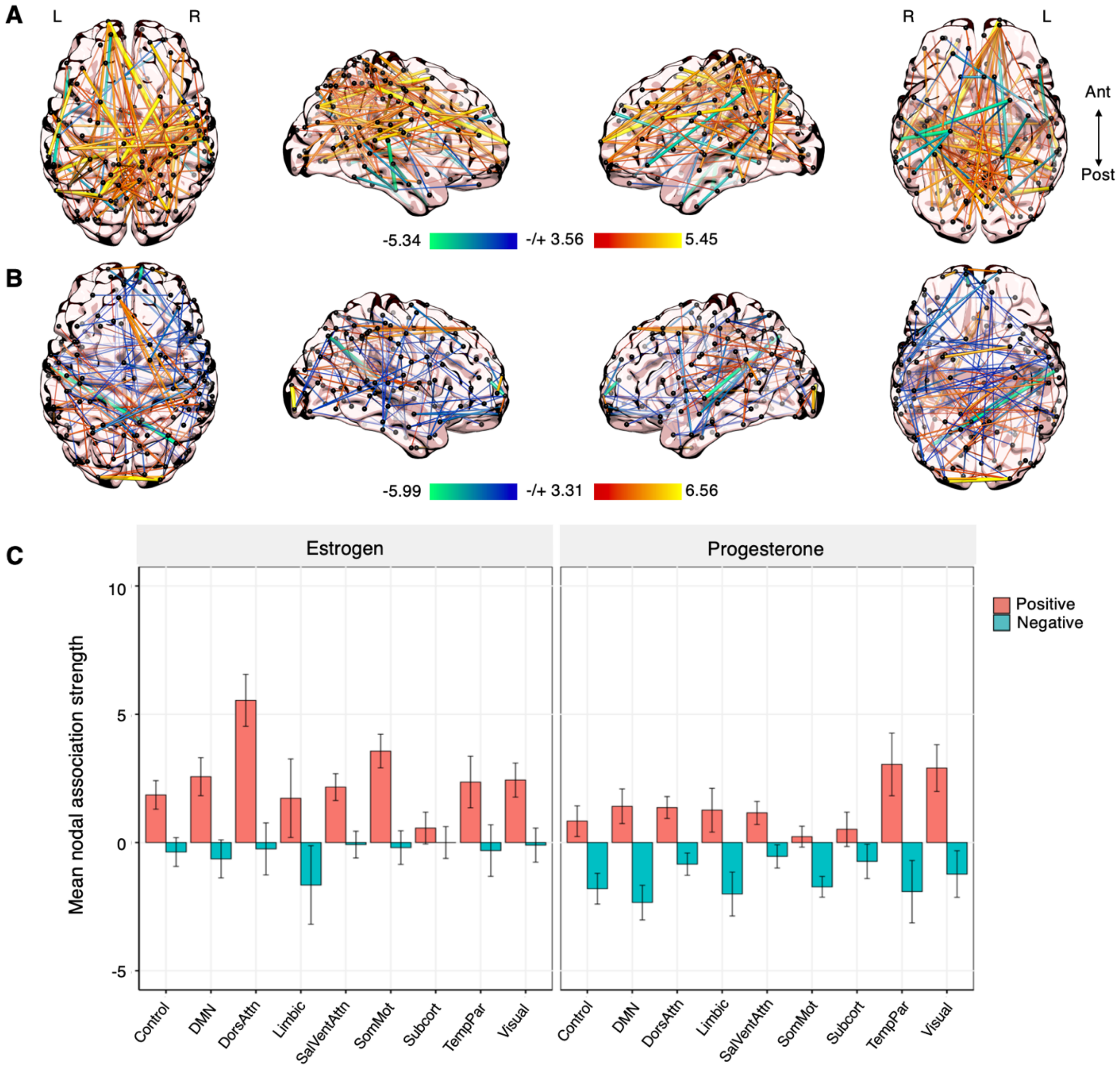


**Figure S5. Global signal regression has minimal impact on time-synchronous brain-hormone associations (Study 1). (A)** Thresholded patterns of association between coherence and estradiol showed moderate degrees of edge-to-edge reliability ($r = 0.48$) after GSR. **(B)** Coherence-progesterone associations after GSR yielded similar degrees of edge-to-edge reliability ($r = 0.61$). **(C)** While the average magnitude of brain-hormone associations differed in some networks, the overall trends remained: namely, that estradiol was associated with widespread increases in coherence.

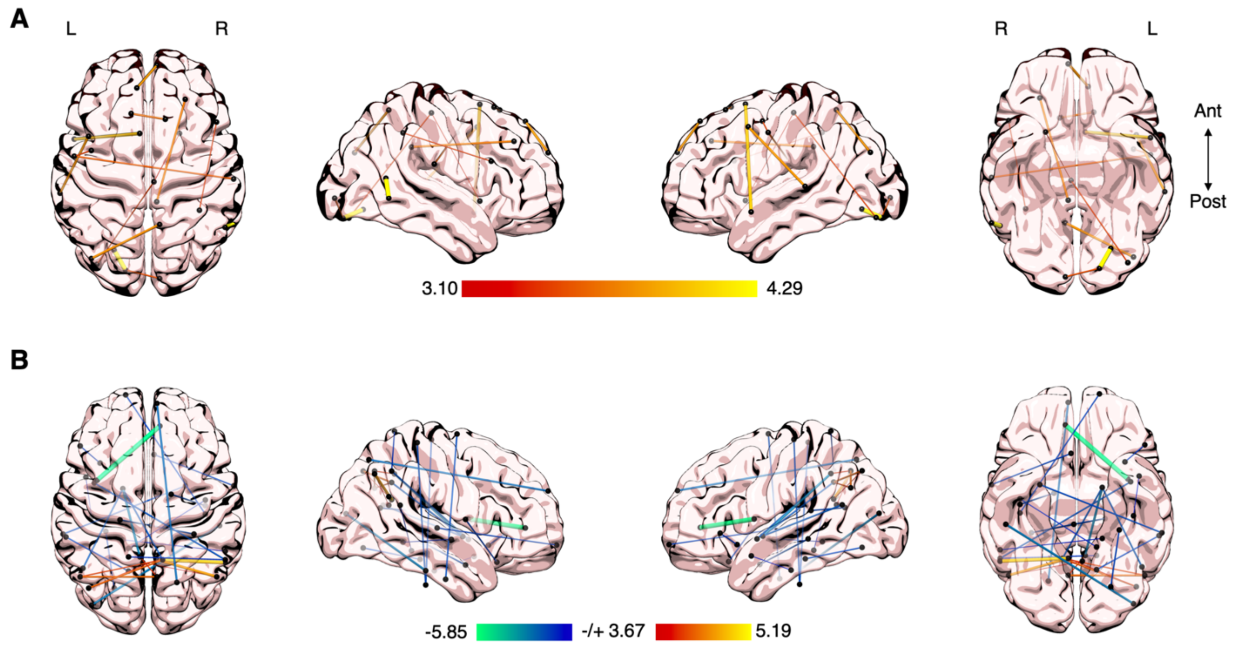


**Figure S6. The ovulatory surge in estradiol is a key modulator of whole-brain functional connectivity. (A)** Removal of the ovulation window in Study 1 (experiment days 22-24) almost entirely erases all significant associations between estradiol and edgewise coherence. **(B)** Removing analogously high estradiol days in Study 2 (experiment days 28-30) also has a strongly-diminishing effect.

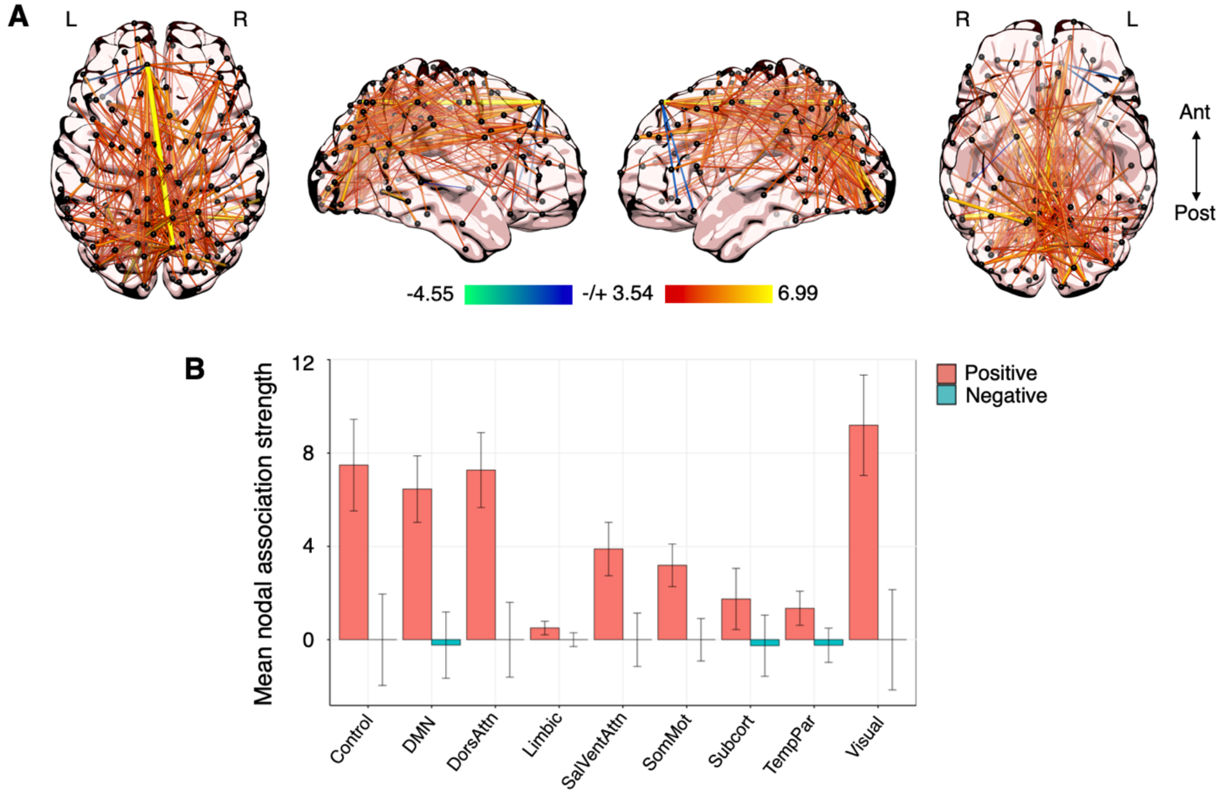


**Figure S7. Whole-brain associations between coherence and estradiol persist in a replication sample (Study 2). (A)** Time-synchronous associations between estradiol and coherence while progesterone was pharmacologically suppressed. Hotter colors indicate increased coherence with higher concentrations of estradiol; cool colors indicate the reverse. Results are empirically-thresholded via 10,000 iterations of nonparametric permutation testing ($p < .001$). Nodes without significant edges are omitted for clarity. **(B)** Mean nodal association strengths by network (with 95% CIs): increased concentrations of estradiol were again coincident with ubiquitous increases in resting connectivity.

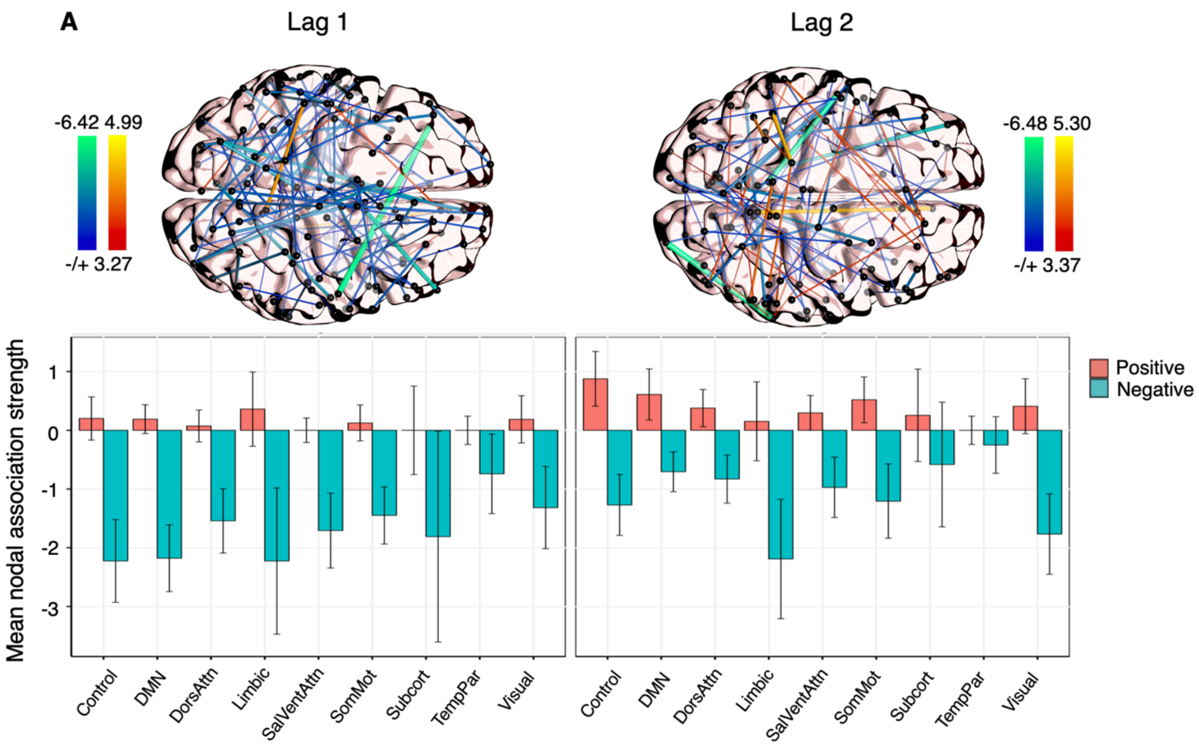


**Figure S8. Autoregressive effects of edgewise coherence (Study 1). (A)** Autoregressive trends in coherence at lag 1 (*left*) and lag 2 (*right*), derived from edgewise vector autoregressive models. Hotter colors indicate a predicted increase in coherence given previous states of connectivity; cool colors indicate the reverse. Results are empirically-thresholded via 10,000 iterations of nonparametric permutation testing ($p < .001$). Nodes without significant edges are omitted for clarity. **(B)** Mean nodal association strengths by network and time lag. Error bars give 95% confidence intervals.

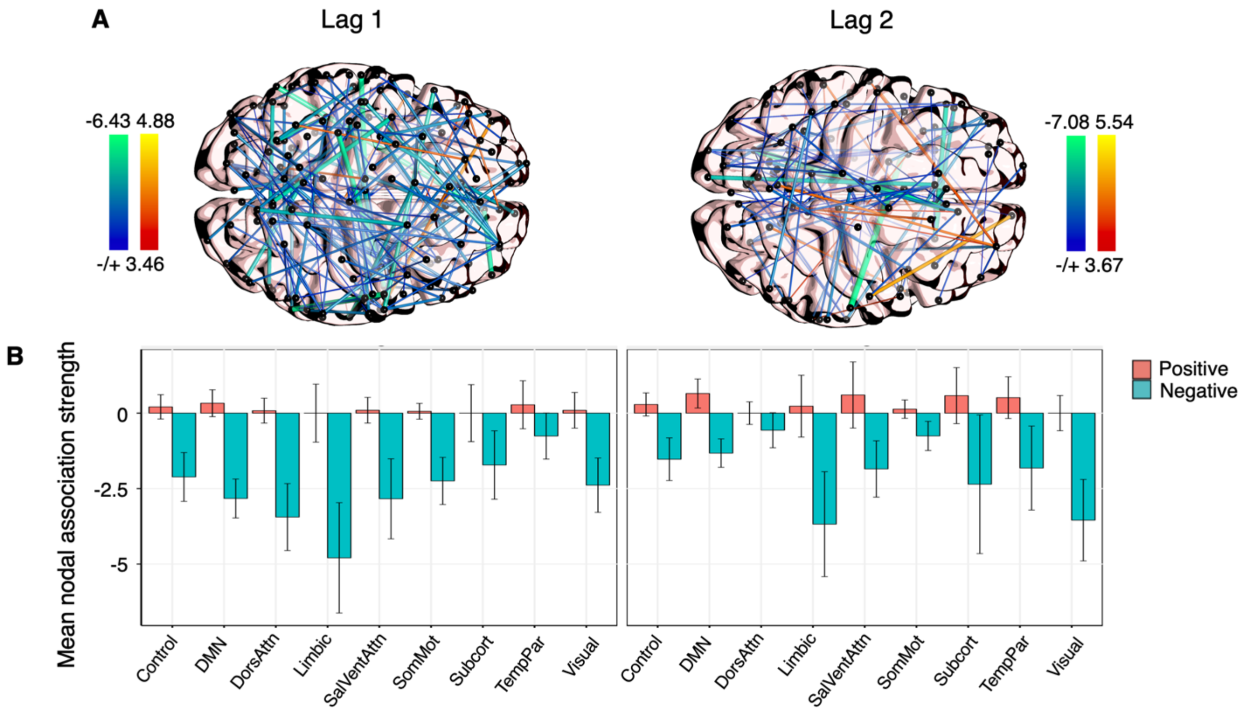


**Figure S9. Estradiol shows linear dependencies on previous states of coherence (Study 1). (A)** Time-lagged associations between estradiol and previous states of coherence at lag 1 (*left*) and lag 2 (*right*), derived from edgewise vector autoregressive models. Hotter colors indicate a predicted increase in estradiol given previous states of coherence; cool colors indicate the reverse. Results are empirically-thresholded via 10,000 iterations of nonparametric permutation testing ($p < .001$). Nodes without significant edges are omitted for clarity. **(B)** Mean nodal association strengths by network and time lag. Error bars give 95% confidence intervals.

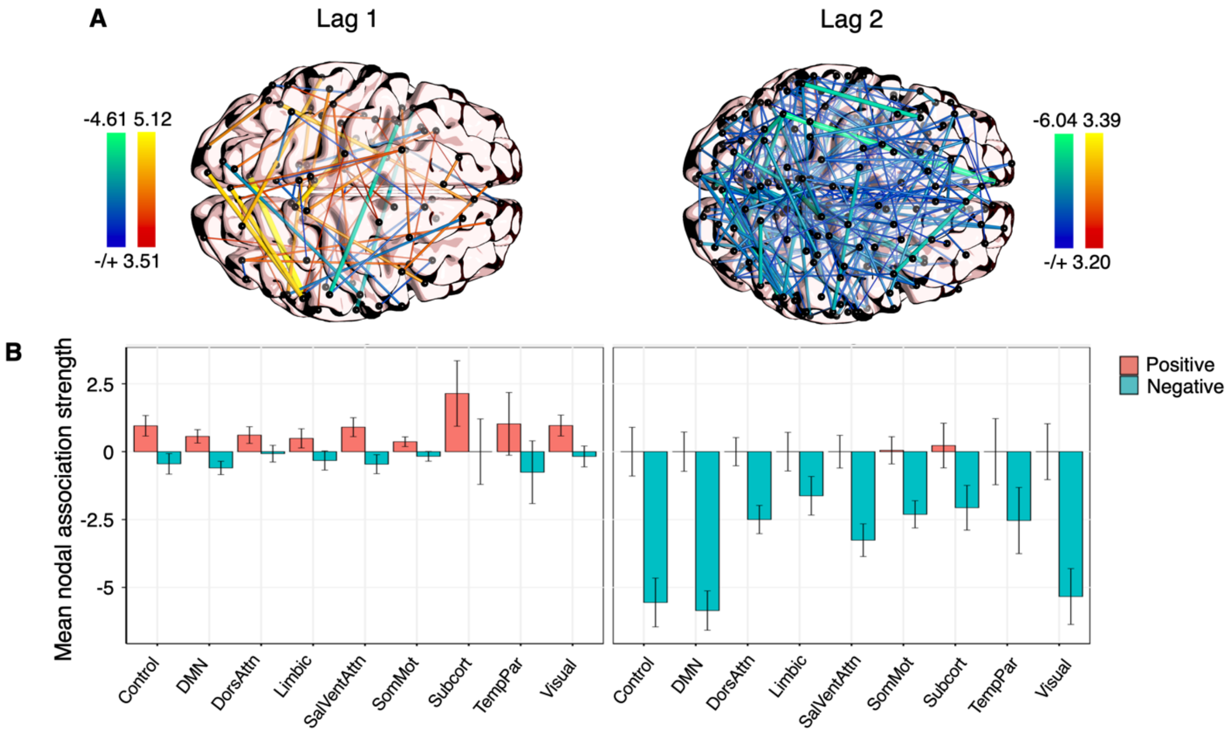


**Figure S10. Autoregressive effects of edgewise coherence (Study 2). (A)** Autoregressive trends in coherence at lag 1 (*left*) and lag 2 (*right*), derived from edgewise vector autoregressive models. Hotter colors indicate a predicted increase in coherence given previous states of connectivity; cool colors indicate the reverse. Results are empirically-thresholded via 10,000 iterations of nonparametric permutation testing ($p < .001$). Nodes without significant edges are omitted for clarity. **(B)** Mean nodal association strengths by network and time lag. Error bars give 95% confidence intervals.

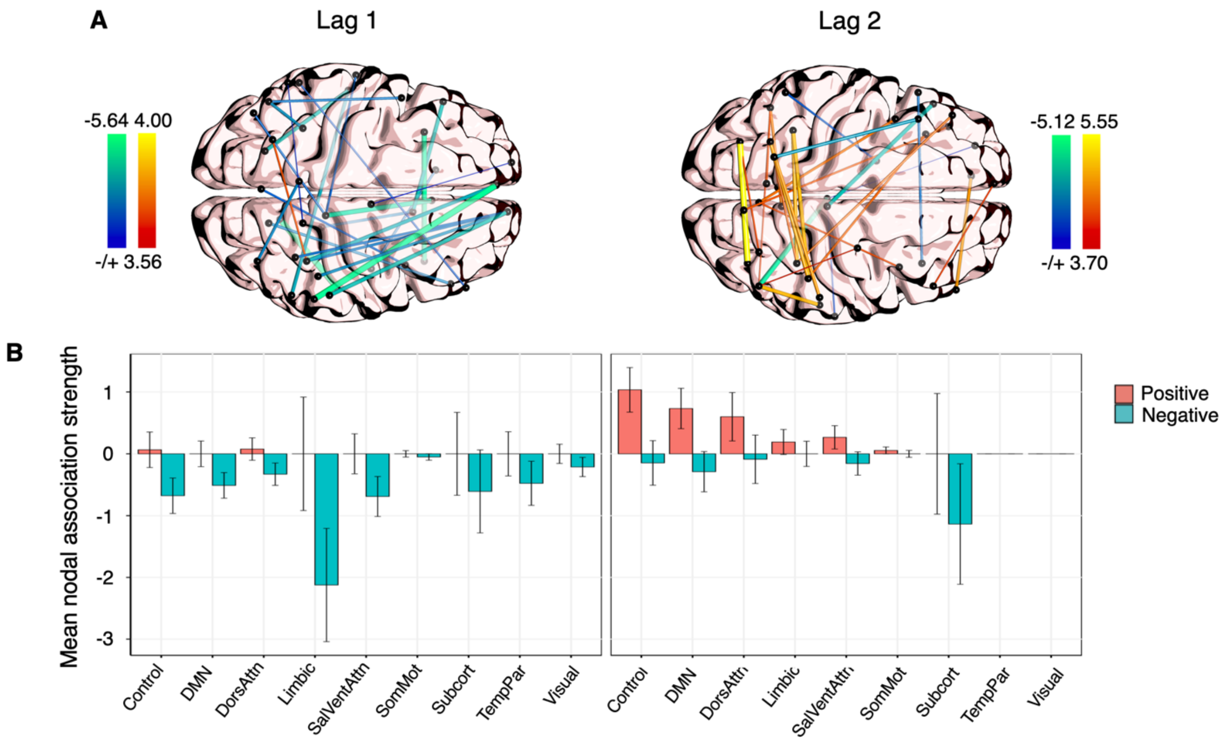


**Figure S11. Estradiol shows linear dependencies on previous states of coherence (Study 2). (A)** Time-lagged associations between estradiol and previous states of coherence at lag 1 (*left*) and lag 2 (*right*), derived from edgewise vector autoregressive models. Hotter colors indicate a predicted increase in estradiol given previous states of coherence; cool colors indicate the reverse. Results are empirically-thresholded via 10,000 iterations of nonparametric permutation testing ($p < .001$). Nodes without significant edges are omitted for clarity. **(B)** Mean nodal association strengths by network and time lag. Error bars give 95% confidence intervals.

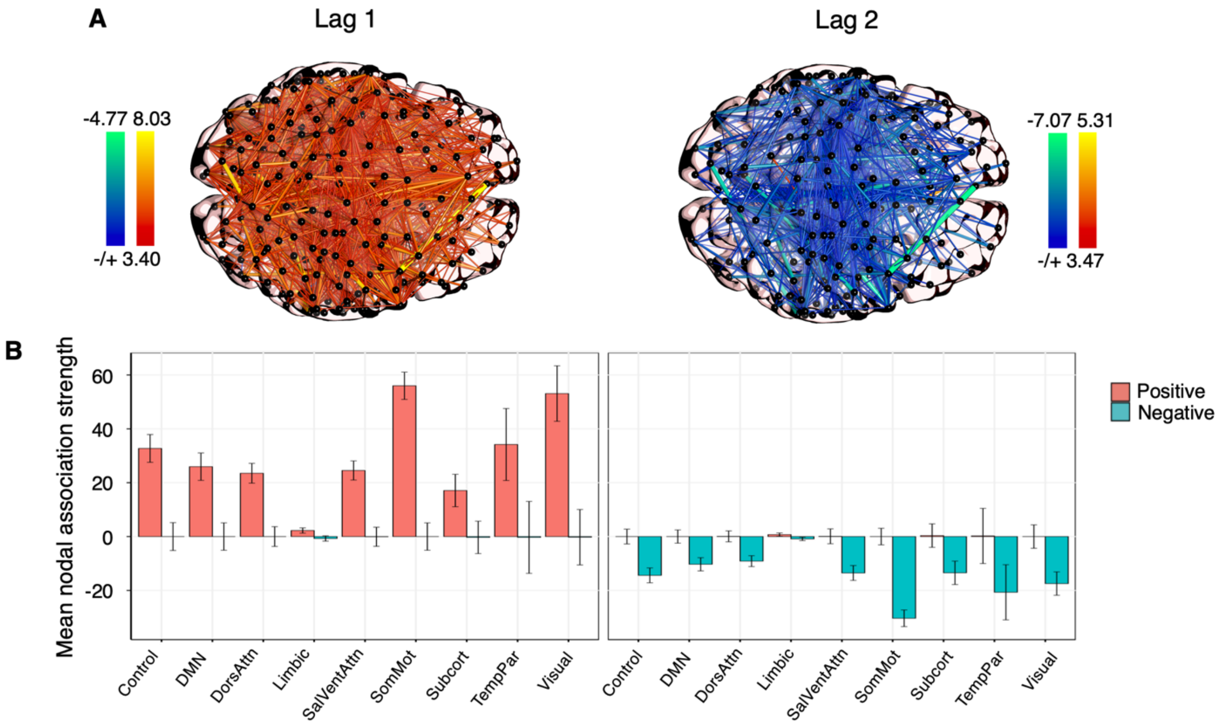


**Figure S12. Linear dependencies between coherence and previous states of estradiol persist in a replication sample (Study 2). (A)** Time-lagged associations between coherence and previous states of estradiol at lag 1 (*left*) and lag 2 (*right*), derived from edgewise vector autoregressive models. Hotter colors indicate a predicted increase in coherence given previous states of estradiol; cool colors indicate the reverse. Results are empirically-thresholded via 10,000 iterations of nonparametric permutation testing ($p <$ .001). Nodes without significant edges are omitted for clarity. **(B)** Mean nodal association strengths by network and time lag. Error bars give 95% confidence intervals. Relative to Study 1, the average magnitude of association by network was greatly increased, perhaps due to the suppression of progesterone.

Table S1. All VAR models for cross-network participation v. estradiol (Study 1).

| **Network** | **Outcome** | **Predictor** | **Estimate** | ***SE*** | ***T* (*p*)** |
|---|---|---|---|---|---|
| *Control* | Participation | **Constant** | **0.11** | **0.18** | **0.62 (.048)** |
| | | $FCN_{t\text{-}1}$ | -0.15 | 0.19 | -0.80 (.430) |
| | | **$Estradiol_{t\text{-}1}$** | **-0.58** | **0.26** | **-2.21 (.036)** |
| | | $FCN_{t\text{-}2}$ | -0.18 | 0.18 | -0.97 (.334) |
| | | $Estradiol_{t\text{-}2}$ | 0.40 | 0.27 | 1.49 (.152) |
| | | | $R^2$ = 0.22 ($p$ = .199); *RMSE* = 0.85 ($p$ = .183) | | |
| | Estradiol | Constant | 0.01 | 0.12 | 0.11 (.691) |
| | | $FCN_{t\text{-}1}$ | -0.04 | 0.13 | -0.27 (.787) |
| | | **$Estradiol_{t\text{-}1}$** | **1.11** | **018** | **6.15 (< .0001)** |
| | | $FCN_{t\text{-}2}$ | -0.03 | 0.13 | -0.27 (.792) |
| | | **$Estradiol_{t\text{-}2}$** | **-0.51** | **0.18** | **-2.75 (.005)** |
| | | | **$R^2$ = 0.66 ($p$ = .0001); *RMSE* = 0.59 ($p$ = .001)** | | |
| *Default Mode* | Participation | Constant | 0.01 | 0.17 | 0.06 (.808) |
| | | $DMN_{t\text{-}1}$ | $-3.13 \times 10^{-4}$ | 0.18 | $-2 \times 10^{-3}$ (.998) |
| | | **$Estradiol_{t\text{-}1}$** | **-0.63** | **0.26** | **-2.43 (.026)** |
| | | $DMN_{t\text{-}2}$ | 0.09 | 0.18 | 0.53 (.599) |
| | | **$Estradiol_{t\text{-}2}$** | **0.72** | **0.23** | **2.84 (.010)** |
| | | | $R^2$ = 0.27 ($p$ = .091); *RMSE* = 0.83 ($p$ = .132) | | |
| | Estradiol | Constant | 0.01 | 0.12 | 0.07 (.803) |
| | | $DMN_{t\text{-}1}$ | -0.08 | 0.12 | -0.66 (.505) |
| | | **$Estradiol_{t\text{-}1}$** | **1.10** | **0.18** | **6.10 (< .0001)** |
| | | $DMN_{t\text{-}2}$ | -0.49 | 0.18 | 0.12 (.906) |
| | | **$Estradiol_{t\text{-}2}$** | **-0.49** | **0.18** | **-2.76 (.004)** |
| | | | **$R^2$ = 0.67 ($p$ < .0001); *RMSE* = 0.57 ($p$ = .0004)** | | |
| *Dorsal Attention* | Participation | Constant | 0.08 | 0.16 | 0.49 (.099) |
| | | $DAN_{t\text{-}1}$ | 0.15 | 0.18 | 0.84 (.405) |
| | | **$Estradiol_{t\text{-}1}$** | **-0.56** | **0.25** | **-2.27 (.035)** |
| | | $DAN_{t\text{-}2}$ | -0.29 | 0.17 | -1.71 (.093) |
| | | **$Estradiol_{t\text{-}2}$** | **0.53** | **0.24** | **2.16 (.042)** |
| | | | **$R^2$ = 0.32 ($p$ = .049); *RMSE* = 0.79 ($p$ = .050)** | | |
| | Estradiol | Constant | $6.88 \times 10^{-5}$ | 0.12 | 0.001 (.998) |
| | | $DAN_{t\text{-}1}$ | 0.06 | 0.14 | 0.47 (.627) |
| | | **$Estradiol_{t\text{-}1}$** | **1.12** | **0.18** | **6.12 (< .0001)** |
| | | $DAN_{t\text{-}2}$ | 0.03 | 0.13 | 0.24 (.806) |
| | | **$Estradiol_{t\text{-}2}$** | **-0.48** | **0.18** | **-2.65 (.007)** |
| | | | **$R^2$ = 0.67 ($p$ = .0001); *RMSE* = 0.59 ($p$ = .0009)** | | |

| | | | | | |
|---|---|---|---|---|---|
| *Limbic* | Participation | Constant | -0.01 | 0.18 | -0.03 (.929) |
| | | $LN_{t\text{-}1}$ | 0.36 | 0.20 | 1.83 (.074) |
| | | **$\mathbf{Estradiol}_{t\text{-}1}$** | **-0.55** | **0.27** | **-2.06 (.050)** |
| | | $LN_{t\text{-}2}$ | -0.09 | 0.20 | -0.45 (.666) |
| | | **$\mathbf{Estradiol}_{t\text{-}2}$** | **0.56** | **0.27** | **2.09 (.047)** |
| | | $R^2$ = 0.27 ($p$ = .102); *RMSE* = 0.86 ($p$ = .235) | | | |
| | Estradiol | Constant | 0.01 | 0.12 | 0.07 (.793) |
| | | $LN_{t\text{-}1}$ | 0.07 | 0.13 | 0.49 (.624) |
| | | **$\mathbf{Estradiol}_{t\text{-}1}$** | **1.13** | **0.18** | **6.34 (< .0001)** |
| | | $LN_{t\text{-}2}$ | -0.14 | 0.13 | -1.05 (.307) |
| | | **$\mathbf{Estradiol}_{t\text{-}2}$** | **-0.52** | **0.18** | **-2.89 (.003)** |
| | | **$R^2$ = 0.68 ($p$ < .0001); *RMSE* = 0.58 ($p$ = .0005)** | | | |
| *Salience/ Ventral Attention* | Participation | Constant | -0.02 | 0.23 | -0.12 (.715) |
| | | $SVN_{t\text{-}1}$ | 0.15 | 0.20 | 0.73 (.488) |
| | | $Estradiol_{t\text{-}1}$ | 0.28 | 0.99 | 0.29 (.782) |
| | | $SVN_{t\text{-}2}$ | 0.11 | 0.22 | 0.51 (.625) |
| | | $Estradiol_{t\text{-}2}$ | -0.12 | 1.18 | -0.10 (.919) |
| | | $R^2$ = 0.07 ($p$ = .80); *RMSE* = 0.96 ($p$ = .87) | | | |
| | Estradiol | Constant | 0.07 | 0.05 | 1.45 (< .0001) |
| | | $SVN_{t\text{-}1}$ | -0.09 | 0.04 | -1.96 (.059) |
| | | **$\mathbf{Estradiol}_{t\text{-}1}$** | **1.31** | **0.21** | **6.27 (< .0001)** |
| | | $SVN_{t\text{-}2}$ | -0.01 | 0.05 | -0.14 (.887) |
| | | $Estradiol_{t\text{-}2}$ | -0.24 | 0.25 | -0.96 (.378) |
| | | **$R^2$ = 0.96 ($p$ < .0001); *RMSE* = 0.20 ($p$ < .0001)** | | | |
| *Somato-Motor* | Participation | Constant | -0.08 | 0.22 | -0.34 (.223) |
| | | $SMN_{t\text{-}1}$ | 0.09 | 0.20 | 0.42 (.664) |
| | | $Estradiol_{t\text{-}1}$ | 0.99 | 0.89 | 1.11 (.270) |
| | | $SMN_{t\text{-}2}$ | 0.17 | 0.20 | 0.84 (.405) |
| | | $Estradiol_{t\text{-}2}$ | -0.87 | 1.07 | -0.82 (.420) |
| | | $R^2$ = 0.13 ($p$ = .46); *RMSE* = 0.92 ($p$ = .50) | | | |
| | Estradiol | Constant | 0.07 | 0.05 | 1.25 (.003) |
| | | $SMN_{t\text{-}1}$ | -0.03 | 0.04 | -0.57 (.568) |
| | | **$\mathbf{Estradiol}_{t\text{-}1}$** | **1.35** | **0.21** | **6.40 (< .0001)** |
| | | $SMN_{t\text{-}2}$ | -0.01 | 0.05 | -0.29 (.767) |
| | | $Estradiol_{t\text{-}2}$ | -0.29 | 0.25 | -1.15 (.266) |
| | | **$R^2$ = 0.96 ($p$ < .0001); *RMSE* = 0.22 ($p$ < .0001)** | | | |

| | | | | | |
|---|---|---|---|---|---|
| *Subcortical* | Participation | Constant | 0.04 | 0.19 | 0.21 (.344) |
| | | SCN$_{t-1}$ | -4.77 x $10^{-3}$ | 0.19 | -0.03 (.976) |
| | | Estradiol$_{t-1}$ | -0.38 | 0.29 | -1.33 (.183) |
| | | SCN$_{t-2}$ | -0.03 | 0.19 | -0.15 (.866) |
| | | Estradiol$_{t-2}$ | 0.58 | 0.28 | 2.04 (.059) |
| | | | | | $R^2$ = 0.16 (*p* = .330); *RMSE* = 0.93 (*p* = .477) |
| | Estradiol | Constant | 0.01 | 0.12 | 0.07 (.792) |
| | | SCN$_{t-1}$ | 0.07 | 0.12 | 0.54 (.573) |
| | | **Estradiol$_{t-1}$** | **1.11** | **0.18** | **6.17 (< .0001)** |
| | | SCN$_{t-2}$ | 0.02 | 0.12 | 0.18 (.854) |
| | | **Estradiol$_{t-2}$** | **-0.50** | **0.18** | **-2.78 (.005)** |
| | | | | | **$R^2$ = 0.67 (*p* < .0001); *RMSE* = 0.59 (*p* = .0009)** |
| *Temporal Parietal* | Participation | Constant | -0.04 | 0.24 | -0.16 (.594) |
| | | TPN$_{t-1}$ | 0.09 | 0.20 | 0.42 (.688) |
| | | Estradiol$_{t-1}$ | 0.27 | 0.97 | .28 (.79) |
| | | TPN$_{t-2}$ | -0.20 | 0.21 | -0.95 (.35) |
| | | Estradiol$_{t-2}$ | -0.34 | 1.15 | -0.28 (.79) |
| | | | | | $R^2$ = 0.05 (*p* = .49); *RMSE* = 0.99 (*p* = .96) |
| | Estradiol | **Constant** | **0.07** | **0.05** | **1.45 (< .0001)** |
| | | TPN$_{t-1}$ | -0.04 | 0.04 | -0.85 (.400) |
| | | **Estradiol$_{t-1}$** | **1.27** | **0.20** | **6.33 (< .0001)** |
| | | TPN$_{t-2}$ | -0.07 | 0.04 | -1.61 (.104) |
| | | Estradiol$_{t-2}$ | -0.22 | 0.24 | -0.93 (.395) |
| | | | | | **$R^2$ = 0.96 (*p* < .0001); *RMSE* = 0.20 (*p* < .0001)** |
| *Visual* | Participation | Constant | 0.17 | 0.22 | 0.62 (.038) |
| | | **VN$_{t-1}$** | **0.09** | **0.21** | **3.27 (.004)** |
| | | Estradiol$_{t-1}$ | 0.03 | 0.89 | -1.57 (.130) |
| | | VN$_{t-2}$ | 0.14 | 0.20 | -1.56 (.140) |
| | | Estradiol$_{t-2}$ | 1.44 | 1.06 | 1.57 (.127) |
| | | | | | **$R^2$ = 0.37 (*p* = .027); *RMSE* = 0.79 (*p* = .05)** |
| | Estradiol | Constant | 0.06 | 0.05 | 1.23 (.003) |
| | | VN$_{t-1}$ | -0.05 | 0.05 | -0.89 (.388) |
| | | **Estradiol$_{t-1}$** | **1.38** | **0.21** | **6.42 (< .001)** |
| | | VN$_{t-2}$ | -0.02 | 0.05 | 0.30 (.776) |
| | | Estradiol$_{t-2}$ | -0.34 | 0.25 | -1.25 (.229) |
| | | | | | **$R^2$ = 0.96 (*p* < .0001); *RMSE* = 0.22 (*p* < .0001)** |

*Note.* *p*-values empirically-derived via 10,000 iterations of nonparametric permutation testing.

Table S2. All re-test VAR models for cross-network participation v. estradiol (Study 2).

| Network | Outcome | Predictor | Estimate | *SE* | *T* (*p*) |
|---|---|---|---|---|---|
| | | Constant | -0.07 | 0.23 | -0.31 (.296) |
| | | $FCN_{t\text{-}1}$ | 0.02 | 0.20 | 0.12 (.905) |
| | Participation | $Estradiol_{t\text{-}1}$ | 0.76 | 0.94 | 0.81 (.429) |
| | | $FCN_{t\text{-}2}$ | -0.18 | 0.20 | -0.87 (.394) |
| | | $Estradiol_{t\text{-}2}$ | -0.75 | 1.12 | -0.67 (.510) |
| | | $R^2$ = 0.08 ($p$ = .739); *RMSE* = 0.96 ($p$ = .834) | | | |
| *Control* | | | | | |
| | | **Constant** | **0.07** | **0.05** | **1.41 (.001)** |
| | | $FCN_{t\text{-}1}$ | -0.05 | 0.04 | -1.07 (.298) |
| | Estradiol | **$Estradiol_{t\text{-}1}$** | **1.31** | **0.20** | **6.61 (< .0001)** |
| | | $FCN_{t\text{-}2}$ | -0.07 | 0.04 | -1.67 (.105) |
| | | $Estradiol_{t\text{-}2}$ | -0.24 | 0.24 | -1.03 (.322) |
| | | **$R^2$ = 0.96 ($p$ < .0001); *RMSE* = 0.20 ($p$ < .0001)** | | | |
| | | Constant | -0.09 | 0.24 | -0.39 (.165) |
| | | $DMN_{t\text{-}1}$ | $-4.00 \times 10^{-3}$ | 0.21 | 0.02 (.985) |
| | Participation | $Estradiol_{t\text{-}1}$ | 0.67 | 0.99 | 0.67 (.510) |
| | | $DMN_{t\text{-}2}$ | -0.12 | 0.21 | -0.54 (.592) |
| | | $Estradiol_{t\text{-}2}$ | -0.73 | 1.18 | -0.62 (.552) |
| | | $R^2$ = 0.05 ($p$ = .891); *RMSE* = 0.99 ($p$ = .937) | | | |
| *Default Mode* | | | | | |
| | | **Constant** | **0.08** | **0.05** | **1.55 (.0002)** |
| | | $DMN_{t\text{-}1}$ | -0.07 | 0.04 | -1.61 (.121) |
| | Estradiol | **$Estradiol_{t\text{-}1}$** | **1.25** | **0.20** | **6.11 (< .0001)** |
| | | $DMN_{t\text{-}2}$ | -0.06 | 0.04 | -1.25 (.214) |
| | | $Estradiol_{t\text{-}2}$ | -0.18 | 0.24 | -0.73 (.503) |
| | | **$R^2$ = 0.96 ($p$ < .0001); *RMSE* = 0.20 ($p$ < .0001)** | | | |
| | | Constant | 0.07 | 0.24 | 0.27 (.347) |
| | | $DAN_{t\text{-}1}$ | $2.00 \times 10^{-3}$ | 0.21 | 0.01 (.991) |
| | Participation | $Estradiol_{t\text{-}1}$ | -0.31 | 1.01 | -0.30 (.766) |
| | | $DAN_{t\text{-}2}$ | 0.01 | 0.23 | 0.05 (.964) |
| | | $Estradiol_{t\text{-}2}$ | 0.61 | 1.19 | 0.51 (.617) |
| | | $R^2$ = 0.04 ($p$ = .909); *RMSE* = 0.99 ($p$ = .988) | | | |
| *Dorsal Attention* | | | | | |
| | | **Constant** | **0.07** | **0.05** | **1.39 (.0007)** |
| | | **$DAN_{t\text{-}1}$** | **-0.09** | **0.04** | **-2.08 (.044)** |
| | Estradiol | **$Estradiol_{t\text{-}1}$** | **1.27** | **0.20** | **6.22 (< .0001)** |
| | | $DAN_{t\text{-}2}$ | -0.04 | 0.05 | -0.85 (.408) |
| | | $Estradiol_{t\text{-}2}$ | -0.21 | 0.24 | -0.88 (.416) |
| | | **$R^2$ = 0.96 ($p$ < .0001); *RMSE* = 0.20 ($p$ < .0001)** | | | |

| | | | | | |
|---|---|---|---|---|---|
| *Limbic* | Participation | **Constant** | **0.15** | **0.21** | **0.70 (.017)** |
| | | **$LN_{t\text{-}1}$** | **0.49** | **0.20** | **2.39 (.029)** |
| | | $Estradiol_{t\text{-}1}$ | -1.27 | 0.84 | -1.52 (.138) |
| | | $LN_{t\text{-}2}$ | -0.28 | 0.19 | -1.47 (.155) |
| | | $Estradiol_{t\text{-}2}$ | 1.54 | 1.01 | 1.53 (.144) |
| | | $R^2$ = 0.24 ($p$ = .17); *RMSE* = 0.85 ($p$ = .160) | | | |
| | Estradiol | **Constant** | **0.07** | **0.05** | **1.27 (.002)** |
| | | $LN_{t\text{-}1}$ | $-7.00 \times 10^{-3}$ | 0.05 | -0.14 (.890) |
| | | **$Estradiol_{t\text{-}1}$** | **1.33** | **0.21** | **6.25 (< .0001)** |
| | | $LN_{t\text{-}2}$ | -0.04 | 0.05 | -0.84 (.406) |
| | | $Estradiol_{t\text{-}2}$ | -0.29 | 0.26 | -1.12 (.282) |
| | | **$R^2$ = 0.96 ($p$ < .0001); *RMSE* = 0.21 ($p$ < .0001)** | | | |
| *Salience/ Ventral Attention* | Participation | Constant | -0.02 | 0.23 | -0.11 (.715) |
| | | $SVN_{t\text{-}1}$ | 0.15 | 0.20 | 0.73 (.488) |
| | | $Estradiol_{t\text{-}1}$ | 0.28 | 0.99 | 0.29 (.782) |
| | | $SVN_{t\text{-}2}$ | 0.11 | 0.22 | 0.51 (.625) |
| | | $Estradiol_{t\text{-}2}$ | -0.12 | 1.18 | -0.10 (.919) |
| | | $R^2$ = 0.07 ($p$ = .802); *RMSE* = 0.96 ($p$ = .874) | | | |
| | Estradiol | **Constant** | **0.07** | **0.05** | **1.45 (.0002)** |
| | | $SVN_{t\text{-}1}$ | -0.09 | 0.04 | -1.96 (.059) |
| | | **$Estradiol_{t\text{-}1}$** | **1.31** | **0.21** | **6.26 (< .0001)** |
| | | $SVN_{t\text{-}2}$ | -0.01 | 0.05 | -0.14 (.887) |
| | | $Estradiol_{t\text{-}2}$ | -0.24 | 0.25 | -0.96 (.378) |
| | | **$R^2$ = 0.96 ($p$ < .0001); *RMSE* = 0.20 ($p$ < .0001)** | | | |
| *Somato-Motor* | Participation | Constant | -0.08 | 0.22 | -0.34 (.224) |
| | | $SMN_{t\text{-}1}$ | 0.09 | 0.20 | 0.42 (.664) |
| | | $Estradiol_{t\text{-}1}$ | 0.99 | 0.89 | 1.11 (.270) |
| | | $SMN_{t\text{-}2}$ | 0.17 | 0.20 | 0.84 (.405) |
| | | $Estradiol_{t\text{-}2}$ | -0.87 | 1.07 | -0.82 (.420) |
| | | $R^2$ = 0.13 ($p$ = .463); *RMSE* = 0.94 ($p$ = .644) | | | |
| | Estradiol | **Constant** | **0.07** | **0.05** | **1.25 (.003)** |
| | | $SMN_{t\text{-}1}$ | -0.03 | 0.05 | -0.57 (.587) |
| | | **$Estradiol_{t\text{-}1}$** | **1.35** | **0.21** | **6.40 (< .0001)** |
| | | $SMN_{t\text{-}2}$ | -0.01 | 0.05 | -0.29 (.767) |
| | | $Estradiol_{t\text{-}2}$ | -0.29 | 0.25 | -1.15 (.266) |
| | | **$R^2$ = 0.96 ($p$ < .0001); *RMSE* = 0.22 ($p$ < .0001)** | | | |

| | | | | | |
|---|---|---|---|---|---|
| *Subcortical* | Participation | **Constant** | **0.12** | **0.19** | **0.62 (.038)** |
| | | **SCN$_{t-1}$** | **0.64** | **0.19** | **3.27 (.004)** |
| | | Estradiol$_{t-1}$ | -1.22 | 0.77 | -1.57 (.130) |
| | | SCN$_{t-2}$ | -0.31 | 0.20 | -1.56 (.140) |
| | | Estradiol$_{t-2}$ | 1.44 | 0.92 | 1.57 (.127) |
| | | **$R^2$ = 0.37 ($p$ = .026); *RMSE* = 0.80 ($p$ = .050)** | | | |
| | Estradiol | **Constant** | **0.06** | **0.05** | **1.23 (.003)** |
| | | SCN$_{t-1}$ | -0.05 | 0.05 | -0.89 (.388) |
| | | **Estradiol$_{t-1}$** | **1.35** | **0.21** | **6.43 (< .0001)** |
| | | SCN$_{t-2}$ | 0.02 | 0.05 | 0.30 (.776) |
| | | Estradiol$_{t-2}$ | -0.31 | 0.25 | -1.25 (.229) |
| | | **$R^2$ = 0.96 ($p$ < .0001); *RMSE* = 0.22 ($p$ < .0001)** | | | |
| *Temporal Parietal* | Participation | Constant | -0.04 | 0.24 | -0.16 (.593) |
| | | TPN$_{t-1}$ | 0.09 | 0.21 | 0.42 (.688) |
| | | Estradiol$_{t-1}$ | 0.27 | 0.97 | 0.28 (.793) |
| | | TPN$_{t-2}$ | -0.20 | 0.21 | -0.95 (.356) |
| | | Estradiol$_{t-2}$ | -0.33 | 1.15 | -0.28 (.787) |
| | | $R^2$ = 0.05 ($p$ = .876); *RMSE* = 0.99 ($p$ = .966) | | | |
| | Estradiol | **Constant** | **-0.04** | **0.05** | **1.45 (.0003)** |
| | | TPN$_{t-1}$ | 1.27 | 0.04 | -0.85 (.404) |
| | | **Estradiol$_{t-1}$** | **-0.07** | **0.20** | **6.33 ( < .0001)** |
| | | TPN$_{t-2}$ | -0.22 | 0.04 | -1.66 (.111) |
| | | Estradiol$_{t-2}$ | 0.07 | 0.24 | -0.93 (.390) |
| | | **$R^2$ = 0.96 ($p$ < .0001); *RMSE* = 0.21 ($p$ < .0001)** | | | |
| *Visual* | Participation | **Constant** | **0.14** | **0.22** | **0.62 (.031)** |
| | | VN$_{t-1}$ | 0.09 | 0.21 | 0.44 (.669) |
| | | Estradiol$_{t-1}$ | 0.03 | 0.89 | 0.03 (.977) |
| | | VN$_{t-2}$ | 0.14 | 0.20 | 0.71 (.485) |
| | | Estradiol$_{t-2}$ | 0.17 | 1.06 | 0.16 (.876) |
| | | **$R^2$ = 0.05 ($p$ = .866); *RMSE* = 0.92 ($p$ = .573)** | | | |
| | Estradiol | **Constant** | **0.06** | **0.05** | **1.22 (.003)** |
| | | VN$_{t-1}$ | -0.05 | 0.05 | -1.11 (.281) |
| | | **Estradiol$_{t-1}$** | **1.38** | **0.21** | **6.70 (<.0001)** |
| | | VN$_{t-2}$ | -0.02 | 0.05 | -0.35 (.718) |
| | | Estradiol$_{t-2}$ | -0.34 | 0.25 | -1.37 (.176) |
| | | **$R^2$ = 0.96 ($p$ < .0001); *RMSE* = 0.21 ($p$ < .0001)** | | | |

*Note. p*-values empirically-derived via 10,000 iterations of nonparametric permutation testing.

Table S3. All VAR models for global efficiency v. estradiol (Study 1).

| Network | Outcome | Predictor | Estimate | *SE* | *T* (*p*) |
|---|---|---|---|---|---|
| | | Constant | -0.02 | 0.17 | -0.14 (.576) |
| | | $FCN_{t-1}$ | -0.13 | 0.18 | -0.73 (.452) |
| | Efficiency | **$Estradiol_{t-1}$** | **0.80** | **0.27** | **2.96 (.007)** |
| | | $FCN_{t-2}$ | -0.12 | 0.19 | -0.66 (.510) |
| | | **$Estradiol_{t-2}$** | **-0.68** | **0.28** | **-2.47 (.023)** |
| | | **$R^2$ = 0.34 ($p$ = .039)**; *RMSE* = 0.83 ($p$ = .133) | | | |
| *Control* | | | | | |
| | | Constant | $3.79 \times 10^{-3}$ | 0.12 | 0.03 (.896) |
| | | $FCN_{t-1}$ | -0.19 | 0.12 | -1.56 (.127) |
| | Estradiol | **$Estradiol_{t-1}$** | **1.17** | **0.18** | **6.41 (< .0001)** |
| | | $FCN_{t-2}$ | $-1.83 \times 10^{-3}$ | 0.13 | -0.04 (.967) |
| | | **$Estradiol_{t-2}$** | **-0.49** | **0.19** | **-2.62 (.007)** |
| | | **$R^2$ = 0.69 ($p$ < .0001); *RMSE* = 0.56 ($p$ < .0001)** | | | |
| | | Constant | 0.04 | 0.15 | 0.28 (.279) |
| | | $DMN_{t-1}$ | -0.04 | 0.16 | -0.27 (.764) |
| | Efficiency | **$Estradiol_{t-1}$** | **0.98** | **0.23** | **3.37 (.0003)** |
| | | $DMN_{t-2}$ | -0.02 | 0.16 | -0.11 (.907) |
| | | **$Estradiol_{t-2}$** | **-0.93** | **0.23** | **-4.00 (.002)** |
| | | **$R^2$ = 0.50 ($p$ = .003); *RMSE* = 0.70 ($p$ = .022)** | | | |
| *Default Mode* | | | | | |
| | | Constant | 0.01 | 0.12 | 0.09 (.729) |
| | | $DMN_{t-1}$ | -0.12 | 0.13 | -0.95 (.339) |
| | Estradiol | **$Estradiol_{t-1}$** | **1.15** | **0.19** | **6.15 (< .0001)** |
| | | $DMN_{t-2}$ | -0.01 | 0.13 | -0.08 (.930) |
| | | **$Estradiol_{t-2}$** | **-0.48** | **0.19** | **-2.50 (.012)** |
| | | **$R^2$ = 0.67 ($p$ < .0001); *RMSE* = 0.58 ($p$ = .0004)** | | | |
| | | Constant | 0.01 | 0.16 | 0.08 (.783) |
| | | $DAN_{t-1}$ | -0.11 | 0.18 | -0.60 (.562) |
| | Efficiency | **$Estradiol_{t-1}$** | **0.84** | **0.25** | **3.35 (.002)** |
| | | $DAN_{t-2}$ | -0.10 | 0.18 | -0.58 (.571) |
| | | **$Estradiol_{t-2}$** | **-0.67** | **0.16** | **-2.57 (.017)** |
| | | **$R^2$ = 0.37 ($p$ = .022); *RMSE* = 0.77 ($p$ = .023)** | | | |
| *Dorsal Attention* | | | | | |
| | | Constant | 0.01 | 0.12 | 0.06 (.808) |
| | | $DAN_{t-1}$ | -0.17 | 0.13 | -1.29 (.207) |
| | Estradiol | **$Estradiol_{t-1}$** | **1.17** | **0.19** | **6.30 (< .0001)** |
| | | $DAN_{t-2}$ | -0.02 | 0.13 | -0.16 (.875) |
| | | **$Estradiol_{t-2}$** | **-0.48** | **0.19** | **-2.49 (.011)** |
| | | **$R^2$ = 0.68 ($p$ < .0001); *RMSE* = 0.57 ($p$ = .0004)** | | | |

| | | | | | |
|---|---|---|---|---|---|
| *Limbic* | Efficiency | Constant | 0.06 | 0.20 | 0.29 (.318) |
| | | $LN_{t\text{-}1}$ | -0.10 | 0.22 | -0.47 (.640) |
| | | $Estradiol_{t\text{-}1}$ | 0.09 | 0.32 | 0.29 (.777) |
| | | $LN_{t\text{-}2}$ | -0.24 | 0.22 | -1.09 (.294) |
| | | $Estradiol_{t\text{-}2}$ | -0.10 | 0.32 | -0.33 (.751) |
| | | | | | $R^2$ = 0.07 (*p* = .760); *RMSE* = 0.97 (*p* = .857) |
| | Estradiol | Constant | 0.03 | 0.11 | 0.31 (.250) |
| | | **$LN_{t\text{-}1}$** | **-0.26** | **0.11** | **-2.26 (.035)** |
| | | **$Estradiol_{t\text{-}1}$** | **1.14** | **0.17** | **6.66 (< .0001)** |
| | | $LN_{t\text{-}2}$ | -0.13 | 0.12 | -1.23 (.235) |
| | | **$Estradiol_{t\text{-}2}$** | **-0.48** | **0.17** | **-2.84 (.004)** |
| | | | | | **$R^2$ = 0.74 (*p* < .0001); *RMSE* = 0.52 (*p* = .0001)** |
| *Salience/ Ventral Attention* | Efficiency | Constant | -0.03 | 0.18 | -0.17 (.533) |
| | | $SVN_{t\text{-}1}$ | -0.05 | 0.19 | -0.27 (.786) |
| | | $Estradiol_{t\text{-}1}$ | 0.45 | 0.28 | 1.58 (.131) |
| | | $SVN_{t\text{-}2}$ | -0.15 | 0.20 | -0.74 (.452) |
| | | $Estradiol_{t\text{-}2}$ | -0.47 | 0.28 | -1.67 (.107) |
| | | | | | $R^2$ = 0.18 (*p* = .309); *RMSE* = 0.88 (*p* = .311) |
| | Estradiol | Constant | -0.01 | 0.11 | -0.07 (.787) |
| | | $SVN_{t\text{-}1}$ | -0.22 | 0.11 | -1.93 (.069) |
| | | **$Estradiol_{t\text{-}1}$** | **1.06** | **0.17** | **6.26 (< .0001)** |
| | | $SVN_{t\text{-}2}$ | -0.18 | 0.12 | -1.57 (.126) |
| | | **$Estradiol_{t\text{-}2}$** | **-0.41** | **0.27** | **-2.41 (.015)** |
| | | | | | **$R^2$ = 0.73 (*p* < .0001); *RMSE* = 0.53 (*p* < .0001)** |
| *Somato-Motor* | Efficiency | Constant | 0.01 | 0.19 | 0.03 (.913) |
| | | $SMN_{t\text{-}1}$ | -0.03 | 0.20 | -0.17 (.856) |
| | | $Estradiol_{t\text{-}1}$ | 0.53 | 0.30 | 1.75 (.093) |
| | | $SMN_{t\text{-}2}$ | -0.07 | 0.22 | -0.30 (.762) |
| | | $Estradiol_{t\text{-}2}$ | -0.47 | 0.31 | -1.52 (.140) |
| | | | | | $R^2$ = 0.15 (*p* = .385); *RMSE* = 0.92 (*p* = .553) |
| | Estradiol | Constant | -0.01 | 0.11 | -0.07 (.775) |
| | | $SMN_{t\text{-}1}$ | -0.21 | 0.11 | -1.89 (.074) |
| | | **$Estradiol_{t\text{-}1}$** | **1.05** | **0.17** | **6.18 (< .0001)** |
| | | $SMN_{t\text{-}2}$ | -0.23 | 0.12 | -1.88 (.076) |
| | | **$Estradiol_{t\text{-}2}$** | **-0.36** | **0.17** | **-2.05 (.036)** |
| | | | | | **$R^2$ = 0.74 (*p* < .0001); *RMSE* = 0.51 (*p* < .0001)** |

| | | | | | |
|---|---|---|---|---|---|
| *Subcortical* | Efficiency | Constant | 0.05 | 0.20 | 0.26 (.351) |
| | | $SCN_{t-1}$ | -0.05 | 0.20 | -0.27 (.788) |
| | | $Estradiol_{t-1}$ | 0.35 | 0.30 | 1.20 (.248) |
| | | $SCN_{t-2}$ | 0.02 | 0.21 | 0.10 (.917) |
| | | $Estradiol_{t-2}$ | -0.38 | 0.30 | -1.29 (.208) |
| | | | | | $R^2$ = 0.07 (*p* = .755); *RMSE* = 0.95 (*p* = .700) |
| | Estradiol | Constant | 0.01 | 0.12 | 0.06 (.823) |
| | | $SCN_{t-1}$ | -0.07 | 0.12 | -0.57 (.597) |
| | | **$Estradiol_{t-1}$** | **1.11** | **0.18** | **6.14 (< .0001)** |
| | | $SCN_{t-2}$ | -0.09 | 0.13 | -0.69 (.493) |
| | | **$Estradiol_{t-2}$** | **-0.47** | **0.18** | **-2.61 (.007)** |
| | | | | | **$R^2$ = 0.67 (*p* < .0001); *RMSE* = 0.58 (*p* = .0006)** |
| *Temporal Parietal* | Efficiency | Constant | 0.01 | 0.16 | 0.90 (.737) |
| | | $TPN_{t-1}$ | -0.21 | 0.18 | -1.15 (.262) |
| | | **$Estradiol_{t-1}$** | **0.89** | **0.26** | **3.40 (.002)** |
| | | $TPN_{t-2}$ | 0.06 | 0.18 | 0.30 (.758) |
| | | **$Estradiol_{t-2}$** | **-0.82** | **0.27** | **-3.04 (.007)** |
| | | | | | **$R^2$ = 0.36 (*p* = .026);** *RMSE* = 0.79 (*p* = .057) |
| | Estradiol | Constant | 0.02 | 0.12 | 0.19 (.477) |
| | | $TPN_{t-1}$ | -0.23 | 0.13 | -1.79 (.087) |
| | | **$Estradiol_{t-1}$** | **1.20** | **0.18** | **6.52 (< .0001)** |
| | | $TPN_{t-2}$ | -0.04 | 0.13 | -0.28 (.783) |
| | | **$Estradiol_{t-2}$** | **-0.50** | **0.19** | **-2.66 (.007)** |
| | | | | | **$R^2$ = 0.70 (*p* < .0001); *RMSE* = 0.55 (*p* = .0005)** |
| *Visual* | Efficiency | Constant | 0.09 | 0.18 | 0.48 (.114) |
| | | $VN_{t-1}$ | -0.10 | 0.18 | -0.54 (.604) |
| | | $Estradiol_{t-1}$ | 0.43 | 0.28 | 1.53 (.142) |
| | | $VN_{t-2}$ | -0.20 | 0.19 | -1.02 (.321) |
| | | $Estradiol_{t-2}$ | -0.37 | 0.28 | -1.32 (.194) |
| | | | | | $R^2$ = 0.18 (*p* = .307); *RMSE* = 0.86 (*p* = .202) |
| | Estradiol | Constant | 0.02 | 0.11 | 0.16 (.534) |
| | | $VN_{t-1}$ | -0.18 | 0.11 | -1.62 (.120) |
| | | **$Estradiol_{t-1}$** | **1.09** | **0.17** | **6.33 (< .0001)** |
| | | $VN_{t-2}$ | -0.19 | 0.12 | -1.63 (.118) |
| | | **$Estradiol_{t-2}$** | **-0.40** | **0.17** | **-2.30 (.019)** |
| | | | | | **$R^2$ = 0.73 (*p* < .0001); *RMSE* = 0.53 (*p* = .0004)** |

*Note. p*-values empirically-derived via 10,000 iterations of nonparametric permutation testing.

Table S4. All re-test VAR models for global efficiency v. estradiol (Study 2).

| Network | Outcome | Predictor | Estimate | *SE* | *T* (*p*) |
|---|---|---|---|---|---|
| | | **Constant** | **-0.12** | **0.21** | **-0.60 (.047)** |
| | | $FCN_{t-1}$ | -0.02 | 0.18 | -0.10 (.921) |
| | Efficiency | **$Estradiol_{t-1}$** | **2.03** | **0.84** | **2.41 (.026)** |
| | | $FCN_{t-2}$ | -0.21 | 0.20 | -1.06 (.292) |
| | | **$Estradiol_{t-2}$** | **-2.14** | **1.02** | **-2.12 (.049)** |
| | | | $R^2$ = 0.23 (*p* = .166); *RMSE* = 0.84 (*p* = .156) | | |
| *Control* | | | | | |
| | | **Constant** | **0.08** | **0.05** | **1.49 (.0003)** |
| | | $FCN_{t-1}$ | 0.03 | 0.05 | 0.73 (.466) |
| | Estradiol | **$Estradiol_{t-1}$** | **1.30** | **0.21** | **6.12 (< .0001)** |
| | | $FCN_{t-2}$ | 0.06 | 0.05 | 1.18 (.253) |
| | | $Estradiol_{t-2}$ | -0.24 | 0.26 | -0.95 (.367) |
| | | | **$R^2$ = 0.96 (*p* < .0001); *RMSE* = 0.21 (*p* < .0001)** | | |
| | | **Constant** | **-0.19** | **0.19** | **-1.04 (.001)** |
| | | $DMN_{t-1}$ | 0.09 | 0.16 | 0.52 (.617) |
| | Efficiency | **$Estradiol_{t-1}$** | **2.48** | **0.75** | **3.29 (.003)** |
| | | **$DMN_{t-2}$** | **-0.45** | **0.19** | **-2.41 (.027)** |
| | | **$Estradiol_{t-2}$** | **-2.69** | **0.91** | **-2.94 (.009)** |
| | | | **$R^2$ = 0.38 (*p* = .019); *RMSE* = 0.74 (*p* = .011)** | | |
| *Default Mode* | | | | | |
| | | **Constant** | **0.08** | **0.05** | **1.44 (.0001)** |
| | | $DMN_{t-1}$ | 0.03 | 0.05 | 0.52 (.598) |
| | Estradiol | **$Estradiol_{t-1}$** | **1.29** | **0.22** | **5.88 (<.0001)** |
| | | $DMN_{t-2}$ | 0.05 | 0.05 | 0.90 (.371) |
| | | $Estradiol_{t-2}$ | -0.24 | 0.27 | -0.90 (.400) |
| | | | **$R^2$ = 0.96 (*p* < .0001); *RMSE* = 0.22 (*p* < .0001)** | | |
| | | Constant | -0.06 | 0.20 | -0.33 (.259) |
| | | $DAN_{t-1}$ | -0.18 | 0.18 | -1.01 (.326) |
| | Efficiency | **$Estradiol_{t-1}$** | **1.88** | **0.79** | **2.37 (.026)** |
| | | $DAN_{t-2}$ | -0.22 | 0.20 | -1.14 (.271) |
| | | $Estradiol_{t-2}$ | -1.64 | 0.94 | -1.74 (.095) |
| | | | $R^2$ = 0.32 (*p* = .052); ***RMSE* = 0.79 (*p* = .045)** | | |
| *Dorsal Attention* | | | | | |
| | | **Constant** | **0.07** | **0.05** | **1.28 (.002)** |
| | | $DAN_{t-1}$ | 0.04 | 0.05 | 0.80 (.442) |
| | Estradiol | **$Estradiol_{t-1}$** | **1.34** | **0.22** | **6.21 (<.0001)** |
| | | $DAN_{t-2}$ | 0.03 | 0.05 | 0.55 (.586) |
| | | $Estradiol_{t-2}$ | -0.31 | 0.26 | -1.20 (.253) |
| | | | **$R^2$ = 0.96 (*p* < .0001); *RMSE* = 0.22 (*p* < .0001)** | | |

| | | | | | |
|---|---|---|---|---|---|
| *Limbic* | Efficiency | Constant | -0.03 | 0.22 | -0.12 (.667) |
| | | $LN_{t-1}$ | 0.38 | 0.20 | 1.91 (.061) |
| | | $Estradiol_{t-1}$ | 0.88 | 0.87 | 1.01 (.327) |
| | | $LN_{t-2}$ | -0.16 | 0.22 | -0.71 (.479) |
| | | $Estradiol_{t-2}$ | -1.05 | 1.06 | -0.99 (.327) |
| | | | | | $R^2$ = 0.15 ($p$ = .389); *RMSE* = 0.88 ($p$ = .274) |
| | Estradiol | **Constant** | **0.07** | **0.05** | **1.23 (.003)** |
| | | $LN_{t-1}$ | 1.00 X $10^{-3}$ | 0.05 | 0.02 (.986) |
| | | **$Estradiol_{t-1}$** | **1.35** | **0.22** | **6.22 (< .0001)** |
| | | $LN_{t-2}$ | 3.00 X $10^{-3}$ | 0.06 | 0.05 (.968) |
| | | $Estradiol_{t-2}$ | -0.30 | 0.27 | -1.14 (.272) |
| | | | | | **$R^2$ = 0.95 ($p$ < .0001); *RMSE* = 0.22 ($p$ < .0001)** |
| *Salience/ Ventral Attention* | Efficiency | **Constant** | **-0.15** | **0.21** | **-0.71 (.017)** |
| | | $SVN_{t-1}$ | 0.14 | 0.18 | 0.75 (.471) |
| | | **$Estradiol_{t-1}$** | **1.98** | **0.87** | **2.27 (.031)** |
| | | $SVN_{t-2}$ | -0.23 | 0.21 | -1.11 (.275) |
| | | **$Estradiol_{t-2}$** | **-2.26** | **1.06** | **-2.14 (.039)** |
| | | | | | $R^2$ = 0.20 ($p$ = .248); *RMSE* = 0.86 ($p$ = .174) |
| | Estradiol | **Constant** | **0.06** | **0.05** | **1.19 (.004)** |
| | | $SVN_{t-1}$ | 0.04 | 0.05 | 0.81 (.416) |
| | | **$Estradiol_{t-1}$** | **1.36** | **0.22** | **6.15 (< .0001)** |
| | | $SVN_{t-2}$ | -1.00 X $10^{-3}$ | 0.05 | -0.01 (.989) |
| | | $Estradiol_{t-2}$ | -0.32 | 0.27 | -1.20 (.245) |
| | | | | | **$R^2$ = 0.95 ($p$ < .0001); *RMSE* = 0.22 ($p$ < .0001)** |
| *Somato-Motor* | Efficiency | **Constant** | **-0.22** | **0.19** | **-1.18 (.0001)** |
| | | $SMN_{t-1}$ | 0.06 | 0.16 | 0.33 (.751) |
| | | **$Estradiol_{t-1}$** | **2.72** | **0.80** | **3.40 (.002)** |
| | | $SMN_{t-2}$ | -0.32 | 0.19 | -1.74 (.097) |
| | | **$Estradiol_{t-2}$** | **-3.16** | **0.97** | **-3.28 (.003)** |
| | | | | | **$R^2$ = 0.34 ($p$ = .039); *RMSE* = 0.76 ($p$ = .018)** |
| | Estradiol | **Constant** | **0.06** | **0.05** | **1.14 (.005)** |
| | | $SMN_{t-1}$ | 0.05 | 0.05 | 1.12 (.268) |
| | | **$Estradiol_{t-1}$** | **1.36** | **0.23** | **6.06 (< .0001)** |
| | | $SMN_{t-2}$ | 4.00 X $10^{-3}$ | 0.05 | 0.07 (.945) |
| | | $Estradiol_{t-2}$ | -0.33 | 0.27 | -1.21 (.236) |
| | | | | | **$R^2$ = 0.96 ($p$ < .0001); *RMSE* = 0.21 ($p$ < .0001)** |

| | | | | | |
|---|---|---|---|---|---|
| | | **Constant** | **-0.19** | **0.21** | **-0.92 (.003)** |
| | | SCN$_{t\text{-}1}$ | 0.22 | 0.18 | 1.24 (.232) |
| | Efficiency | **Estradiol$_{t\text{-}1}$** | **1.89** | **0.83** | **2.29 (.032)** |
| | | SCN$_{t\text{-}2}$ | -0.24 | 0.19 | -1.29 (.204) |
| | | **Estradiol$_{t\text{-}2}$** | **-2.22** | **1.00** | **-2.23 (.037)** |
| | | | | | $R^2$ = 0.23 (*p* = .172); *RMSE* = 0.85 (*p* = .159) |
| *Subcortical* | | | | | |
| | | **Constant** | **0.07** | **0.05** | **1.23 (.002)** |
| | | SCN$_{t\text{-}1}$ | 0.03 | 0.05 | 0.61 (.550) |
| | Estradiol | **Estradiol$_{t\text{-}1}$** | **1.37** | **0.22** | **6.41 (< .0001)** |
| | | SCN$_{t\text{-}2}$ | -0.01 | 0.05 | -0.13 (.901) |
| | | Estradiol$_{t\text{-}2}$ | -0.32 | 0.26 | -1.26 (.216) |
| | | | | | **$R^2$ = 0.95 (*p* < .0001); *RMSE* = 0.22 (*p* < .0001)** |
| | | **Constant** | **-0.17** | **0.22** | **-0.77 (.010)** |
| | | TPN$_{t\text{-}1}$ | 0.16 | 0.20 | 0.83 (.412) |
| | Efficiency | Estradiol$_{t\text{-}1}$ | 1.82 | 0.93 | 1.95 (.059) |
| | | TPN$_{t\text{-}2}$ | -0.31 | 0.21 | -1.42 (.168) |
| | | Estradiol$_{t\text{-}2}$ | -2.28 | 1.14 | -2.00 (.057) |
| | | | | | $R^2$ = 0.16 (*p* = .360); *RMSE* = 0.88 (*p* = .286) |
| *Temporal Parietal* | | | | | |
| | | **Constant** | **0.06** | **0.05** | **1.17 (.004)** |
| | | TPN$_{t\text{-}1}$ | 0.04 | 0.05 | 0.74 (.465) |
| | Estradiol | **Estradiol$_{t\text{-}1}$** | **1.38** | **0.23** | **6.00 (< .0001)** |
| | | TPN$_{t\text{-}2}$ | 0.01 | 0.05 | 0.25 (.797) |
| | | Estradiol$_{t\text{-}2}$ | -0.33 | 0.28 | -1.18 (.257) |
| | | | | | **$R^2$ = 0.95 (*p* < .0001); *RMSE* = 0.22 (*p* < .0001)** |
| | | Constant | -0.11 | 0.21 | -0.50 (.080) |
| | | VN$_{t\text{-}1}$ | -0.05 | 0.19 | -0.27 (.792) |
| | Efficiency | Estradiol$_{t\text{-}1}$ | 1.34 | 0.84 | 1.59 (.126) |
| | | VN$_{t\text{-}2}$ | -0.09 | 0.19 | -0.45 (.659) |
| | | Estradiol$_{t\text{-}2}$ | -1.27 | 1.01 | -1.26 (.227) |
| | | | | | $R^2$ = 0.15 (*p* = .431); *RMSE* = 0.88 (*p* = .264) |
| *Visual* | | | | | |
| | | **Constant** | **0.06** | **0.05** | **1.20 (.003)** |
| | | VN$_{t\text{-}1}$ | 0.02 | 0.05 | 0.41 (.695) |
| | Estradiol | **Estradiol$_{t\text{-}1}$** | **1.38** | **0.21** | **6.60 (< .0001)** |
| | | VN$_{t\text{-}2}$ | -0.03 | 0.05 | -0.69 (.499) |
| | | Estradiol$_{t\text{-}2}$ | -0.33 | 0.25 | -1.34 (.183) |
| | | | | | **$R^2$ = 0.95 (*p* < .0001); *RMSE* = 0.22 (*p* < .0001)** |

*Note. p*-values empirically-derived via 10,000 iterations of nonparametric permutation testing.
